# Phosphorylation sites are evolutionary checkpoints against liquid-solid transition in protein condensates

**DOI:** 10.1101/2022.09.15.508150

**Authors:** Srivastav Ranganathan, Pouria Dasmeh, Seth Furniss, Eugene Shakhnovich

## Abstract

Assemblies of multivalent RNA-binding protein FUS can exist in the functional liquid-like state as well as less dynamic and potentially toxic amyloid- and hydrogel-like states. How could then cells form liquid-like condensates while avoiding their transformation to amyloids? Here we show how post-translational phosphorylation can provide a “handle” that prevents liquid-solid transition of intracellular condensates containing FUS. Using residue-specific coarse-grained simulations, for 85 different mammalian FUS sequences, we show how the number of phosphorylation sites and their spatial arrangement affect intracluster dynamics preventing conversion to amyloids. All atom simulations further confirm that phosphorylation can effectively reduce the β-sheet propensity in amyloid-prone fragments of FUS. A detailed evolutionary analysis shows that mammalian FUS PLDs are enriched in amyloid-prone stretches compared to control neutrally evolved sequences suggesting that mammalian FUS proteins evolved to self-assemble. However, in stark contrast to proteins that do not phase-separate for their function, mammalian sequences have phosphosites in close proximity to these amyloid-prone regions. These results suggest that evolution uses amyloid-prone sequences in prion-like domains to enhance phase-separation of condensate proteins while enriching phosphorylation sites in close proximity to safe-guard against liquid-solid transitions.

**Significance Statement:** Intrinsically disordered regions and prion-like domains are widely observed in proteins that get enriched in membrane-less organelles (MLOs). Mammalian Fused in Sarcoma (FUS) sequences are significantly enriched in amyloid-prone sequences suggesting that they have evolved to self-assemble. While the amyloid-prone stretches promote self-assembly of these proteins at lower threshold concentrations, these assemblies are vulnerable to aberrant liquid-solid phase transitions. Molecular simulations and bioinformatics analyses show that evolution overcomes this challenge by placing phosphosites specifically close to amyloid-prone stretches. Introduction of negatively charged residues at phosphosite locations results in fewer amyloid-prone contacts and thereby lower beta-sheet propensity. Phosphorylation can thus allow cells to utilize amyloid-prone stretches to promote biogenesis of MLOs while protecting against liquid-solid transitions.

## Introduction

Biomolecular phase transitions is a universal mechanism of formation of “membraneless organelles” (MLOs) in mammalian cells. These organelles have been associated with diverse functions ranging from stress response to gene regulation ^1–4^. A candidae physical mechanism to explain the formation of MLOs is intracellular liquid-liquid phase separation (LLPS) wherein inter-molecular interactions result in formation of a biopolymer-rich phase that is distinct from the bulk solution ^1,5^. A key feature of proteins that engage in intracellular condensates is multivalency – the ability to participate in several adhesive interactions^6,7^.

Multivalency can take different forms, including folded domains capable of specific protein-protein interactions in modular proteins, or low-complexity, intrinsically disordered regions that can participate in multiple attractive inter-protein interactions^1,3^. Over the past decade, intrinsically disordered regions (IDRs) have been identified to be widely prevalent in phase-separating proteins^8,9^. The multivalent interactions involving IDRs has been found to reduce threshold concentrations for phase-separation of folded domains by an order of magnitude^10^. While the multivalency of IDRs stems from their ability to participate in diverse interactions (pi-pi/cation, hydrophobic, electrostatics) and thus allows LLPS to occur at physiological concentrations, a high density of prion-like domains (PLDs) within the condensed phase makes these structures prone to detrimental, liquid-solid phase transitions. Indeed, PLDs of RNA-binding domains have been observed to populate diverse physical states such as liquid-droplets, hydrogels and ordered amyloid-like structures^11–13^. This suggests an interesting paradox. While the multivalency of IDRs stems from their ability to participate in diverse interactions (pi-pi/cation, hydrophobic, electrostatics) and thus allows LLPS to occur at physiological concentrations, a high density of prion-like domains (PLDs) within the condensed phase makes these structures prone to detrimental, liquid-solid phase transitions. Full-length FUS proteins that initially self-assemble into dynamic, liquid-like droplets over time evolve into amyloid-like solid structures in a process known as maturation of droplets in vitro^13^. These results suggest that the amyloid-like state could potentially be an equilibrium configuration for IDR-rich phase-separated structures. How then cells strike a balance between the functional role of IDRs (ability to phase separate) and their deleterious effects (irreversible liquid-solid transitions) is an open question.

Active mechanisms such as post-translational modifications have been suggested to play an important role in preventing liquid-solid transitions in the cell^14^. Phosphorylation, methylation, acetylation and ribosylation have all been reported in intrinsically disordered regions of phase-separating proteins^15^. Previous studies, both *in vitro* and *in silico* have suggested the ability of phosphorylation to prevent fibrillation in FUS droplets^16–20^. Monahan et al. ^19^ employed phosphomimics to show that phosphorylation disrupts aggregation in vitro and shifts threshold concentration for assembly of FUS LC domain. Interestingly, while 32 putative phosphorylation sites have been identified in the FUS PLD (residues 1-165), the FUS protein doesn’t always get extensively phosphorylated in response to DNA damage^15,18^. Identification of partially phosphorylated forms of FUS under some conditions suggests that cells could exploit differential phosphorylation patterns for fine-grain control over the fate of FUS assemblies^15,18^. The biological relevance of stochasticity in phosphorylation patterns is further emphasized in studies where only a small fraction of FUS shows complete phosphorylation^18,21^. The nature of the physical stress has also been shown to impact phosphorylation pattern, with Ser-42 phosphorylation being a more prevalent modification in response to IR damage, while UV radiation resulted in an increase in Ser-26 modification^18,21^. The computational study by Perdikari et al^20^ shows variability in FUS conformations in a phosphorylation pattern dependent manner. What are the mechanistic origins of this phosphorylation pattern-dependent variability? Is the effect of phosphorylation simply net-charge driven, or does evolution select for phosphosites at specific locations along the FUS PLD sequence? While earlier studies point out the important role of phosphorylation as a potential modulator of phase-separation, the mechanistic and evolutionary origins of phosphorylation as a tunable handle for FUS self-assembly remain largely unaddressed.

In this study we employ a residue-specific coarse-grained model for FUS as well as all atom simulations and evolutionary analysis to study how phosphorylation influences the inter-protein interaction network as well as intracluster dynamics in phase-separated FUS proteins. Using a residue-specific coarse-grained model proposed by Dignon et al^22^, we show how intracluster dynamics of FUS PLD assemblies is primarily mediated by Tyrosine and Glutamine residues. We further study the effect of phosphorylation on phase behavior of condensates by systematically varying the extent of phosphorylation as well as generating different phosphorylation patterns. Our results show that phosphorylation results in a switch in the predominant interaction network to heterodomain interactions, for full-length FUS assemblies. Importantly, we show that the intracluster dynamics, and diffusion coefficients are not only sensitive to the net charge of the FUS prion-like domain but also the location of the modification sites. All atom simulations further support the role of phosphorylation in drastically diminishing the propensity of amyloid-rich fragments of FUS sequences to form aberrant beta structure. Using simulations and bioinformatics analysis with 85-different mammalian FUS sequences, we also show that the number, and location of phosphosites are strongly correlated with the overall amyloidigenic propensity of a sequence, and, more specifically phosphorylation sites appear to be preferentially located near amyloid-rich segments of FUS proteins. Crucially, proteins that are not known to phase separate for their function do not show such an enrichment of phosphosites. Overall, our study shows that phosphorylation sites in IDRs likely serve as evolutionary checkpoints against solidification via amyloid-like transitions.

## Model

### Coarse-grained flexible polymer model for FUS

We performed Langevin dynamics simulations using a flexible polymer representation of FUS wherein we model the protein at residue level details. We employ a coarse-grained framework previously developed by Dignon et al. for simulating phase-separation of proteins such as LAF1 and the low complexity domain of FUS^22^. Here, each amino-acid is modeled as a single bead in a bead-spring polymer with the physicochemical properties of the bead (hydrophobicity, charge) being mapped onto one of 20 naturally occuring amino acids. To model inter- and intra-chain protein-protein interactions, we employ a hydrophobicity-based interaction potential used by Dignon et al. to study LAF1 phase separation^22^, and first described by Ashbaugh and Hatch^23^. This approach, refered to as the hydrophobicity scale (HPS) method, uses the hydrophobicity scale to define interacions betweena amino acids. The hydrophobicity values for the coarse-grained amino acid beads are scaled from 0 to 1 (with 1 being the most hydrophobic amino acid.) The net hydrophobic strength of an interaction between two coarse-grained amino acids is the arithmetic mean of their individual hydrophobicities (Supplementary Table.1). The Ashbaugh-Hatch potential describing interaction between coarse-grained amino acids assumes the form^22^,

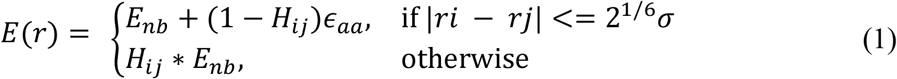

and

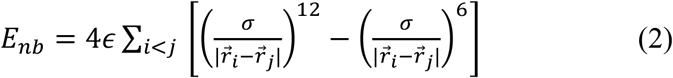

for all 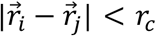. Here *r*_*c*_ refers to the interaction cutoff beyond which LJ interactions are neglected. The cutoff for LJ calculations were set to 2.5 times of *σ*, the size of the interacting beads. In supplementary Table S1, we list the bead sizes for the 20 amino acid bead types used in our simulations. Here, H refers to the net hydrophobicity of an interacting amino-acid pair which is the arithmetic mean of their individual hydrophobicities while *ϵ*_*aa*_ is the strength of the attractive interaction which gets scaled depending on the properties of the amino acid. *ϵ*_*aa*_ was set to 0.2 kcal/mol, a value which resulted in experimentally comparable values for radius of gyration values of intrinsically disordered proteins in the study by Dignon et al^22^. In order to model phosphomimetic FUS sequences, we substitute the phosphosite S/T residues to Glutamic acid (S/T -> E), introducing a negative charge at the site. Similar approach to study phosphorylation of the FUS prion-like domain has been adopted by Perdikari et al^20^ in their simulation study and Monahan et al in vitro^19^. It must, be noted that there are other higher resolution coarse-grained models for simulating intrinsically disordered proteins, notably the work by Baul et al which efficiently captures structural heterogeneity in proteins such as alpha-synuclein, osteopontin and nucleoporin^24^. The choice of the HPS model for our study was dictated by its previous use for specifically studying the phase-separation behavior of FUS making it an ideal reference point for comparisons for our current work. It would, however, be an interesting and useful exercise in a future study to understand how a switch to a more elaborate model (such as the Baul et al.^24^ model which captures SAXS data) influences simulation results.

## Methods

### Coarse-grained simulations

Peptides/proteins have been extensively modeled at coarse-grained resolution to study biomolecular structure and self-assembly^6,25–28^. Fewer degrees of freedom in coarse-grained polymers leads to higher computational tractability, making them efficient tools to study biomolecular phase transitions^27,29–34^. In order to study the effect of phosphorylation on LLPS of FUS and the resultant intra-cluster dynamics, we performed Langevin dynamics simulations. To simulate self-assembly, we perform simulations with 50 coarse-grained FUS (full-length or PLD) chains in a cubic box with periodic boundaries enabled.

Coarse-grained FUS polymer beads interact via the following bonded and non-bonded potential functions. Adjacent beads in a polymer chain are bonded via the harmonic potential with energy

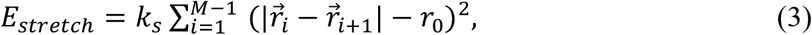

where 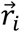 and 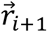 correspond to the positions of the *i*^*th*^ and (*i* + 1)^*th*^ beads, respectively; *r*_0_ is the resting bond length. *k*_*s*_ denotes the spring constant and has units of kT/Å^2^. This interaction ensures the connectivity between the adjacent beads of the same polymer chain. For the coarse-grained FUS polymers, we use a *k*_*s*_ of 2 kT//Å^2^.

To model the flexible nature of the self-assembling intrinsically disordered proteins, neighboring bonds in a polymer chain interact via a bending potential

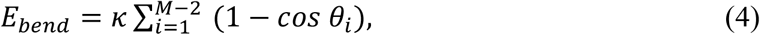

where *θ*_*i*_ refers to the angle between *i*^*th*^ and (*i* + 1)^*th*^ bond. Here, *κ* the bending stiffness, has units energy, kT. We employ a *κ* of 4 kT or equivalently, a persistence length of roughly 5 polymer beads (or 5 amino acid residues). The contour length of the FUS prion-like domain, in comparision is 165 residues long.

Pairwise non-bonded interactions were modeled using a combination of Lennard-jones and Ashbaugh and Hatch potential (see Model section for details). Screened electrostatic interactions were modeled using the Debye-Huckel electrostatic potential,

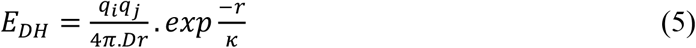

Here, *q*_*i*_ and *q*_*j*_ are the charges of the two coarse-grained amino acid beads (Table S1). This approach has been previously employed in phase separation simulations by Dignon et al.^20,25,35^and can recapitulate behavior of intrinsically disordered proteins such as TDP43 and FUS. The Debye-Huckel screening length of 1 nm was employed in our coarse-grained simulations to mimic the screening of electrostatic interactions in a cellular environment. The above potential energy functions (eqns 1,2 and 5) are used to model both inter- and intra-chain interactions.

The LAMMPS molecular dynamics package was employed to perform the coarse-grained Langevin dynamics simulations^36^. Here, the simulator solves Newton’s equations in presence of a viscous force while the Langevin thermostat is used to implement the NVT ensemble with a simulation temperature *of* 310 K. An integration timestep (*dt*) of 15 fs and a damping time of 3000 fs were used for the simulations (please also see the LAMMPS simulation script in Supplementary Information for reference). The damping time for the thermostat was set to 3500 fs.

### Simulation protocol – coarse-grained simulations

To ensure efficient sampling of the self-assembly landscape, and to access a single large spherical condensate at simulation timescales, we employed metadynamics simulations^37^. In these history-dependent metadynamics simulations, energy gaussians are deposited during the course of the simulation along a reaction coordinate that describes self-assembly. Here, the reaction coordinate along which the metadynamics simulation is performed is the radius of gyration of an imaginary polymer -- R_g_^system^ -- that is composed of the 100 center of masses corresponding to the constituent polymer chains in the system. In Supplementary Fig.S1, we show the free energy profile of a system of 100 FUS chains at an effective concentration of 100 uM. Values of R^g^_system_ approaching 150 Angstroms corresponds to a single large spherical condensate while at higher values (> 200 Angstroms), the system approaches the fully mixed state. As evident from the free-energy profile, the system of FUS proteins favors the demixed, self-assembled state.

These metadynamics simulations enable us to sample the equilibrium, single-large cluster at simulation timescales. This configuration corresponding to the dense phase (the free energy minima in metadynamics simulations) was then used as a starting configuration for conventional Langevin dynamics trajectories (with no biasing potential) of at least 10 microseconds or till the simulations reached steady state. The steady state section of these unbiased 10-microsecond long simulation trajectories were used to compute statistical quantities (inter-protein contacts, diffusion coefficients) in this paper. The order parameters reported in this work are thus independent of the choice of metadynamics collective variable.

### Modeling folded domains in coarse-grained simulations of full-length FUS

While the full-length FUS protein is predominantly disordered, there are regions in the RNA-binding domain – the RNA recognition motif (RRM)-and the zinc-finger motif (ZnF)-that are folded in structure. In our coarse-grained simulations involving the full-length FUS protein, we used the experimentally resolved solution structures of the RRM (PDB ID:6GBM) and ZnF (PDB ID: 6G99). These RRM region (residues 285-371) and the ZnF motif (residues 422-453) were modeled as rigid bodies (with no intrinsic dynamics) using the “fix rigid” command in LAMMPS. The rest of the protein with significant disorder are treated as flexible polymers. There are no interactions within residues of the same rigid body. Inter-chain interactions involving residues of the folded domains, on the other hand, are set to 40% weaker than that of those involving disordered regions to model the solvent-inaccesible nature of these residues. These assumptions are consistent with the treatment of folded regions in LAF1 in a previous study by Dignon et al.^20,35^

### All-atom molecular dynamics

To study secondary structural propensity of amyloidogenic peptides derived from the FUS prion-like domain (PLD), we employed atomistic molecular dynamics simulations. To study the β-sheet propensity of the FUS PLD-derived peptides and the role of phosphorylation in abrogating inter-peptide β-sheet formation, we performed metadynamics simulations with 50 molecules of peptides derived from FUS_1-165_ and their phosphomimetic counterparts (see Supplementary Table. S4 for sequences). The radius of gyration of the system of 50 peptides (R_g_^system^) was used as a reaction coordinate along which the metadynamics^37^ simulations were performed. The structures corresponding to the free energy minima in metadynamics simulations were used to analyze secondary structural propensity of the peptides. The simulations were performed using the NAMD^38^ molecular dynamics package with an electrostatic cutoff of 12 Å units and a van der Waals cutoff of 10 Å units along with periodic boundary conditions. The systems were first energy-minimized and then heated to 310 K with a gradual scaled increase in temperature. This was followed by a constant pressure equilibration for 5 ns and then an NPT (isothermal-isobaric ensemble) production run. The CHARMM36 protein forcefield^39^, and the TIP3P water model was used to simulate proteins at atomistic resolution. All simulations were performed until the simulations reached steady state.

### Simulation of neutrally evolved sequences

We employed the Evolver package within the PAML suite^40^ to simulate protein evolution and to generate neutrally evolved sequences. This approach uses Monte Carlo simulations to generate codon sequences using a specified phylogenetic tree with given branch lengths, nucleotide frequencies, transition/transversion bias, and the ratio of the rate of nonsynonymous to synonymous substitutions (dN/dS; ^41^). For neutral sequence evolution, we used the standard genetic code with codon frequencies taken from the sequence of our proteins of interest. We performed two sets of simulations. In the first set and to compare human proteins with neutrally-evolved sequences, we generated a single protein with the length of 4000 codons. In the second set, and in the analysis of 85 mammalian FUS sequences, we generated 100 neutrally-evolved FUS sequences each with the length of 600 codons. We reported our measured quantities per 100 amino acids to account for different lengths of mammalian PLD sequences and our simulated sequences. We used a consensus phylogenetic tree for mammals from the TimeTree database (^42^). In our phylogenetic trees, the branch lengths represent the expected number of nucleotide substitutions per codon. To model neutral evolution, we set d*N*/d*S* to 1 and estimated transition/transversion rate ratio from the sequence of each of our human proteins and the orthologous protein sequence in Chimpanzee. This ratio was 4.0 for FUS, 2.6 for Osteopontin, 3.2 for LAF-1, 2.4 for Securin, and 4.2 for RYBP. We estimated these values by fitting the codon model M1 to the phylogenetic tree and the sequences of these proteins^43^. This model assumes that all branches of the phylogenetic tree have the same rate of evolution.

### Nomenclature

The 526-residue long FUS protein will be referred to as the full-length FUS. The FUS prion-like domain, on the other hand, will hereby be referred to as FUS_1-165_.

## Results

### Tyrosine and Glutamine contacts are modulators of intracluster dynamics

The FUS family of proteins, including the FUS itself, feature a multi-domain architecture with a low complexity prion-like domain, an RNA-recognition motif and an arginine rich C-terminal domain^12^]. Previous studies have extensively established the different interaction networks and the hierarchy of interactions stabilizing the liquid-like FUS condensate^12^. The PLDs of FUS are also known to self-assemble into hydrogels and solid-like structures^11^. Given the association between interaction networks and the material state of the condensate, it becomes critical to understand the interactions stabilizing the condensed phase.

To identify the key ‘sticky’ interactions in the PLD, we performed simulations with 165-residue long coarse-grained peptide chains (same as the length of the PLD) of the following sequences - i) *polyG*_165_, ii) *polyY*_165_, iii) *polyST*_165_, iv) FUS_1-165_, v) FUS_1-165_ with Glycines replaced by Glutamines (G->Q), vi) FUS_1-165_ with glycines substituted with serine (G->S) and vii) sequences where serine or glutamine residues are replaced by glycines (S>G and Q>G). These substitutions were performed on the 4 most abundant residue types in FUS_1-165_ -- G/Q/S/Y (Fig.1A). Using the coarse-grained forcefield for phase-separating proteins first developed by Dignon et al ^35^, we first perform metadynamics (see methods) simulations to enable the formation of the condensed phase at simulation timescales. We subsequently run long trajectories (10 μs) to analyze the interaction networks within protein clusters. As shown in Fig.1B, the polyG chains remain in a monomeric state and develop negligible interpeptide contacts. polyY, chains on the other hand show the strongest interpeptide contact development within the condensed phase. polyST sequences form self-assembled structures with sparser inter-molecular interactions than polyY chains. The native PLD sequence, on the other hand, shows an intermediate level of inter-peptide contacts, sparser than polyY chains and stronger than polyST chains. Interestingly, modifying the glycine residues in the FUS PLD to glutamine and serine does’nt result in a significant increase in interpeptide contacts. However, mutating the glycine residues of the FUS PLD to either Glutamine or Serine (G->S and G->Q in Fig.1C) results in a dramatic slowdown in intracluster environments. Conversely, modifying S/Q -> G results in assemblies that are more dynamic than the FUS_1-165_ assemblies (Fig. 1B). Interestingly, G->S and S->G substitutions are the most frequent substitutions among mammalian FUS orthologs^44^. These results show that while Tyrosine residues are the strongest drivers of phase-separation in the FUS PLD (Fig. 1A), the intracluster dynamics within the dense phase depends significantly on inter-peptide contacts involving Tyr and Gln (purple and black curves in Fig.1C). These results are consistent with experiments and simulations exploring FUS interaction network^12,13,22^ suggesting that the coarse-grained model can robustly capture features of FUS phase separation. Simulations with homopolymer and amino-acid substituted sequences also suggest that interpeptide contacts involving Tyr and Gln could be robust predictors of intra-cluster dynamics.

**Figure 1.**
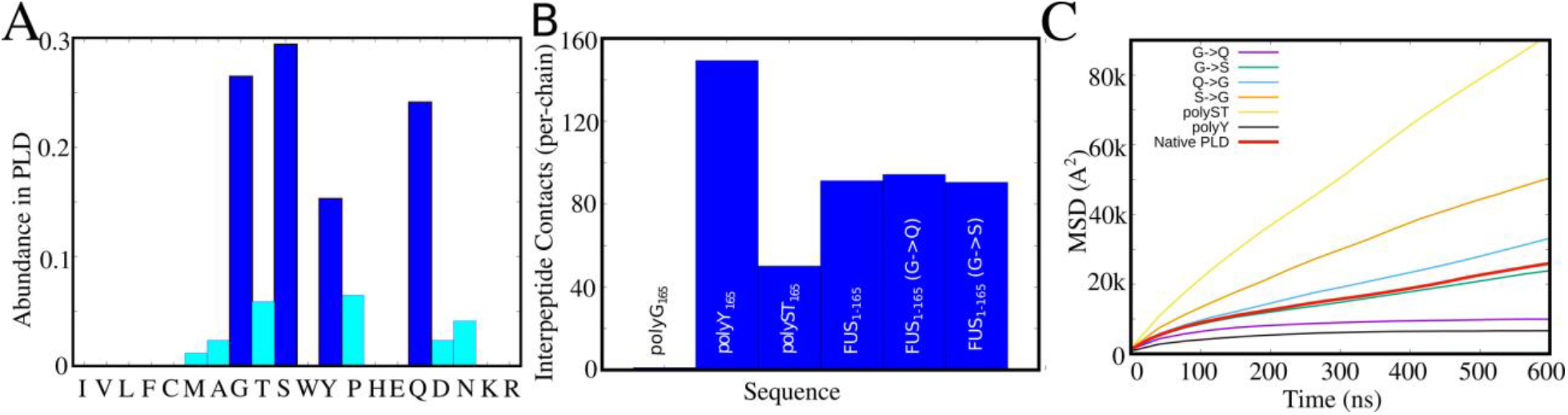
Primary Modulators of Droplet Dynamics. A) Amino acid residues and their abundances in the FUS PLD B) Comparison of inter-peptide contacts (per chain) for different 165-residue long peptide chains. *polyG*_165_ chains remain in the monomeric state. polyY chains show the highest inter-peptide interaction propensity while polyST sequences show an intermediate degree of interaction. The native FUS PLD clusters show inter-peptide contacts sparser than polyY clusters but greater than polyST clusters. C) The mean-square displacement profiles for different 165-residue long peptides. polyST sequences form self-assembled clusters that are more liquid-like than polyY and native FUS PLD clusters. polyY clusters are the most solid-like among all sequences studied. Mutations to glycine residues (G->S and G->Q) results in a slowdown in intracluster dynamics. On the other hand, mutating S/Q -> G results in a more dynamic intracluster environment than the native FUS PLD sequence.

Upon identifying how interactions involving the 4 most-abundant amino acids in the FUS PLD can modulate the intracluster dynamics within protein assemblies, we study whether (and how) shuffling of the PLD sequence in full-length FUS protein could tune the interaction network within the FUS condensates. To understand the key contacts that stabilize the assembly, we randomize the sequence of the FUS PLD (residues 1-165), keeping the amino acid composition of the PLD intact (Supplementary Fig.S2A) and the RNA-binding domain (RBD) sequence unchanged. We observed that shuffling of the PLD sequence results in an almost 2-fold variation in the number of PLD-PLD contacts within the condensed phase (Supplementary Fig.S2B). Crucially, the number of PLD-PLD contacts in shuffled sequences is strongly correlated with number of interactions involving Tyrosine and Glutamine residues in the PLD (Supplementary Fig.S2C and D). However, an increase in Tyr and Gln contacts does not lead to an increase in contacts involving Ser and Gly (Supplementary Fig.S3), the other two most abundant residues in the PLD of FUS (Supplementary Fig.2A), suggesting that the density of the PLD-PLD interactions is primarily modulated by the Tyrosine and Glutamine residues.

### Phosphorylation reduces the likelihood of PLD-PLD contacts while the PLD-RBD interactions remain unaffected

The coarse-grained model can capture the known features of FUS interaction networks where Gly/Ser are spacers and Tyr/Gln are stickers thereby making it suitable to study the role of phosphorylation on the FUS condensate interaction network **(Fig 1 and S1)**. In order to model phosphorylated FUS sequences, we replace the S/T residues at phosphosites to Glutamic acid (S/T -> E), thereby introducing a negative charge at the site. The human FUS PLD sequence has 31 potential phosphosites^45^ (Supplementary Table S2 and S3). We systematically vary the fraction of phosphosites **(P**_**frac**_) that are modified, and also simulated different phosphorylation patterns for each modification fraction. As we increase the fraction of phosphosites that are modified -- **(P**_**frac**_**) --**, we observe an increased likelihood of the heterodomain interactions (Fig. 2A). For **P**_**frac**_ ⟶ **0**, the PLDs are equally likely to participate in homodomain (with other PLDs) or heterodomain interactions (with RBDs of other molecules). On the other hand, as **P**_**frac**_ ⟶ **0.9**, we see a 2-fold higher likelihood of PLD-RBD interactions as compared to the PLD-PLD interactions.

**Figure 2:**
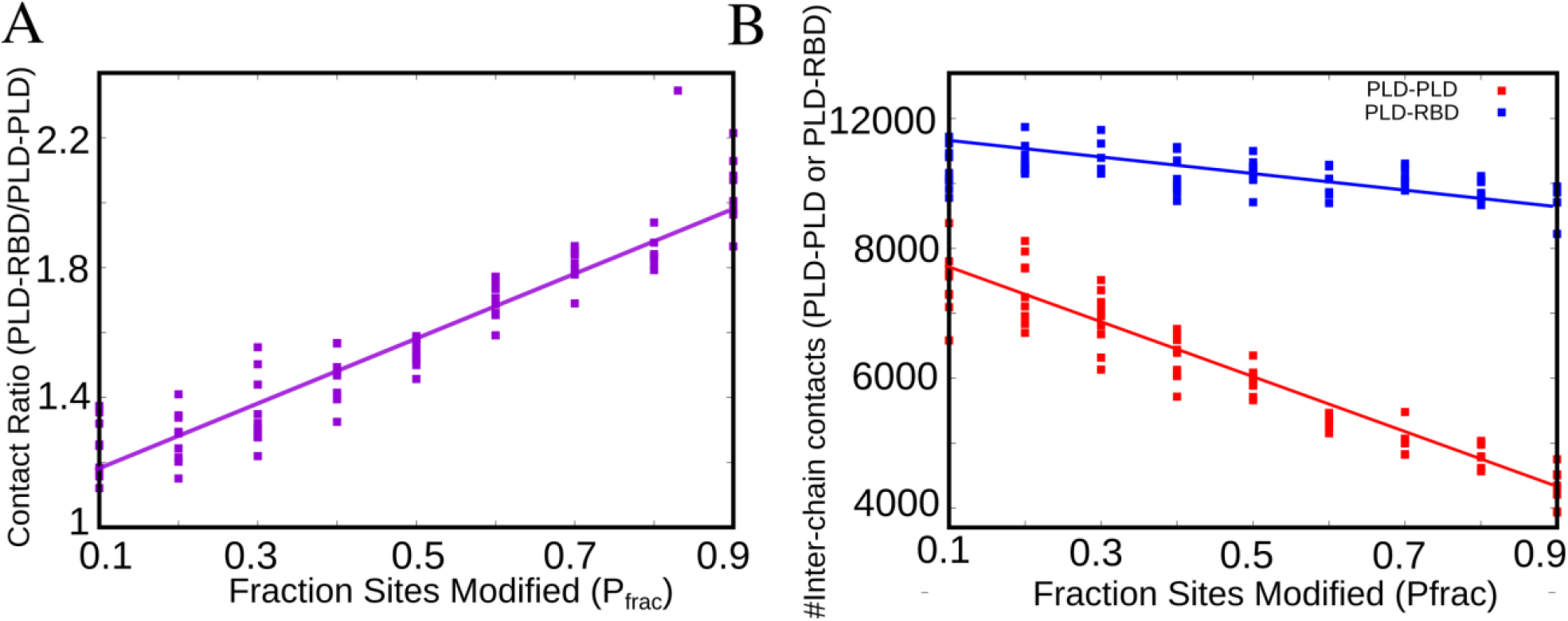
Network switching in full-length FUS as a result of Phosphorylation. A) The ratio of inter-peptide PLD-RBD to PLD-PLD contacts within the self-assembled cluster as a function of varying modification fraction– fraction of residues modified out of a potential 31 phosphosites. B) The effect of phosphorylation on PLD-PLD and PLD-RBD interaction network. Each point in the figure corresponds to a different modification pattern, for a given fraction of sites modified.

Further, we probed whether this switch in interaction networks is an outcome of reduced PLD-PLD interactions or increased PLD-RBD interactions. We find that an increase in modification fraction does not significantly alter the extent of interchain PLD-RBD interactions (Fig 2B, blue curve). On the other hand, the PLD-PLD interactions show a 2-fold decrease (Fig 2B, red curve) as we increase the modification fraction from 0.1 to 0.9 suggesting that phosphorylation can affect interaction networks by selectively modulating PLD-PLD interactions.

### Phosphorylation maintains liquid-like intracluster environment for PLD assemblies

The FUS PLD (FUS_1-165_) has been observed to self-assemble into liquid-like droplets and undergo liquid-to-solid transitions in vitro^19,46^. Also, previous coarse-grained simulations have shown that disorder-order and liquid-solid transitions in FUS are characterized by homotypic contacts involving the PLD^28^. Our simulations with phosphorylated full-length FUS sequences establish that the introduction of negative charges in the PLD selectively tunes PLD-PLD interactions (Fig.2B). Therefore, the effect of phosphorylation can be effectively understood by studying FUS PLDs alone. Hereon, we switch to simulations of PLDs (FUS_1-165_ instead of the full-length protein), allowing for increased computational tractability of the simulations.

We first systematically vary the fraction of modified phosphosites, **(P**_**frac**_), with different phosphorylation patterns for each fraction, and study the effect on the inter-molecular PLD-PLD contacts (Fig.3A). As with the full length FUS, we observe a decrease in PLD-PLD interactions at higher **P**_**frac**_. However, unlike the full-length protein, at **P**_**frac**_ > 0.6 the PLD-peptides remain in monomeric state (Fig.3A, black curve and Supplementary Fig.S4). This is consistent with experiments which show that phosphorylation results in an increase in threshold concentrations for phase separation of PLDs ^47^. The full-length protein, on the other hand, continues to remain in the phase-separated form even at high modification fractions, due to stabilization by heterodomain interactions (Fig.3A, purple curve). In Supplementary Fig.S4, we plot the size of the largest cluster size (L_clus_) as a function of (P_frac_). While full-length FUS remains in the self-assembled state throughout (L_clus_ -> 1), at P_frac_ > 0.6, the largest cluster size for FUS_1-165_ << 1. Hence, even at high modification fractions, the inter-peptide interactions for full-length FUS do not vanish unlike the PLD-only simulations with FUS_1-165 chains_ (Supplementary Fig. S4) due to the presence of the second interaction network – PLD-RBD – in the full-length protein assemblies. This is consistent with previous experimental findings by Murthy et al^48^ as well as Wang et al^12^ which suggests that the N-terminal low-complexity domain interacts with the RNA-binding domain via pi-cation interactions.

**Figure 3:**
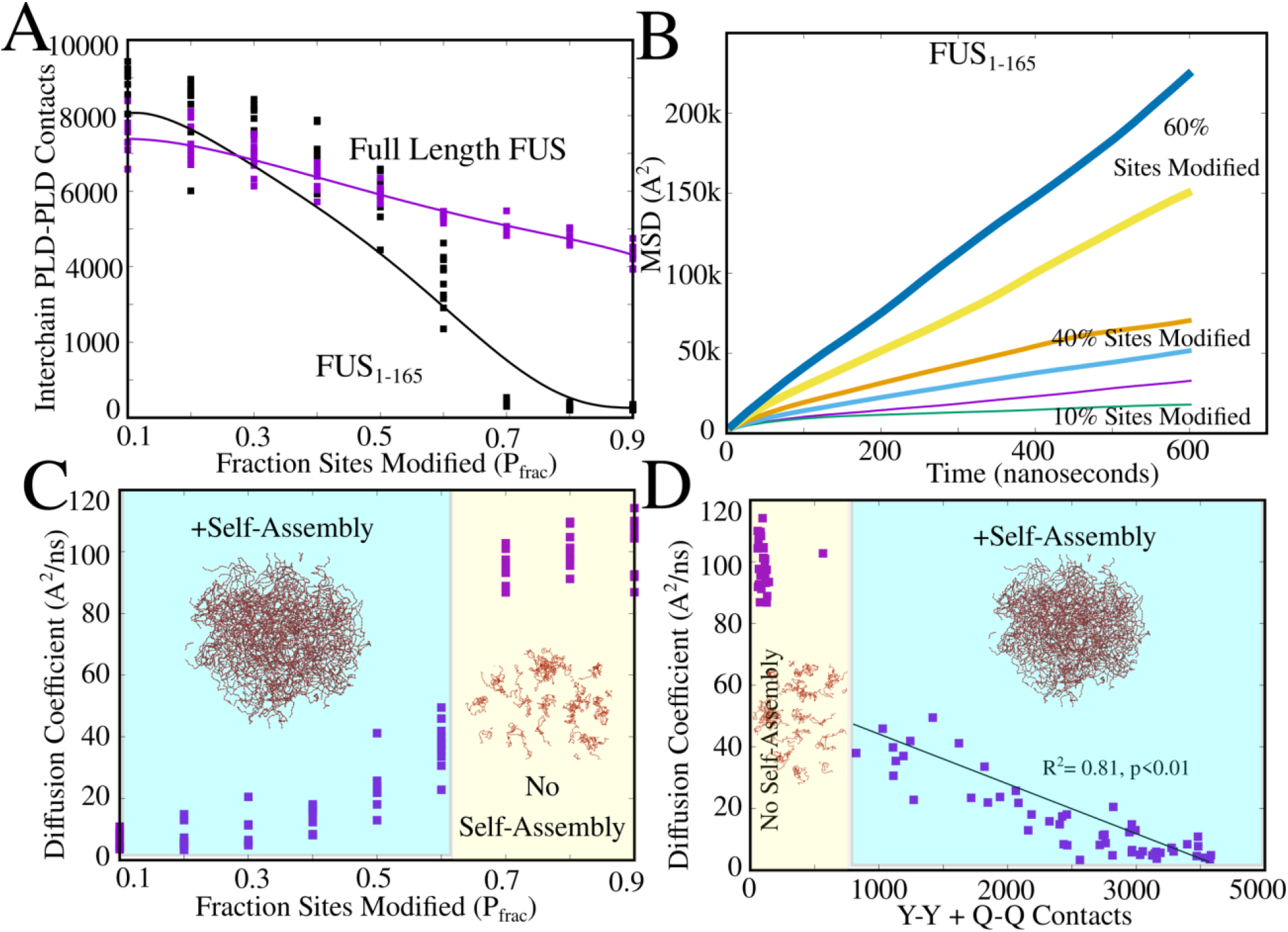
Phosphorylation tunes intracluster dynamics in FUS_1-165_ assemblies. A) PLD-PLD contacts as a function of increasing fraction of phosphosites modified. Purple curve corresponds to full-length FUS (PLD+RBD) while black curve shows results from FUS_1-165_ simulations. Each point corresponds to a different modification pattern, for any **P**_**frac**_. B) Representative mean-square displacement profiles for polymer chains within the FUS_1-165_ self-assembly. Higher the slope of the curve, more dynamic the intra-droplet environment. C) Diffusion coefficients within the FUS_1-165_ assembly (obtained by fitting MSD profiles to *MSD* = 6 * *D* * t. D) Diffusion coefficients are inversely correlated to the number of sticky interactions (Tyr-Tyr and Gln-Gln) within a cluster. +Self-assembly refers to the regime where the largest cluster sizes, L_clus_⟶1, a single large cluster. In the - Self-assembly regime (shaded yellow), L_clus_<<1 (see Supplementary Fig.S4 for cluster sizes).

In Supplementary Fig.S5, we plot the dense and dilute phase concentrations of protein as a function of temperature. For a lower P_frac_ of 0.1, the dense phase remains stable for a larger range of temperatures. For highly phosphorylated sequences (P_frac_ = 0.7), the dense phase is destabilized at critical temperatures below 310K, the temperature at which all the results in this paper are reported. The bulk concentrations at which we observe self-assembled structures of FUS_1-165_ was close to 0.5 mg/ml which is in the range of previously reported threshold concentrations for FUS^35^. The concentration of FUS in the dense phase, on the other hand, was two orders of magnitude higher than that in the bulk.

We further probe whether this reduced amount of PLD-PLD interactions at higher P_frac_ has an effect on intracluster dynamics of proteins. In Fig.3B, we plot the mean square displacement of polymer chains within the self-assembled cluster as a function of time. As we increase the extent of phosphorylation, the slope of MSD curves progressively increases suggesting more-liquid like behavior of phosphorylated PLDs. Interestingly, even for the same P_frac_, we observe variability in MSD profiles (and inter-chain contacts) for different modification patterns. We further compute the diffusion exponents from the mean-square displacement profiles (*MSD* = 6*Dt*). The diffusion coefficients corresponding to different modification fractions and patterns is shown in Fig. 3C. At lower P_frac,_ the clusters are more likely to exhibit slow intracluster dynamics, as evident from the smaller values of the diffusion constants. As we increase the extent of phosphorylation, we see a concommitant increase in the diffusion coefficients suggesting more dynamic intra-cluster environment for these sequences. The diffusion coefficient is strongly correlated with the number of sticky contacts involving Tyr and Gln within the cluster (Fig.3B), consistent with experimental findings that show that stronger involvement of glutamines in inter-peptide interactions slows down intra-cluster dynamics^12^. Varying the extent of phosphorylation results in a 4-fold variation in diffusion coefficients within the self-assembled state (see region shaded blue in Fig. 3C). Overall, these findings suggest that the extent of phosphorylation and its specific patterns can not only alter the threshold concentrations for phase-separation (Fig.3A and Supplementary Fig. S4,S5) but also serve as effective modulator of intra-condensate dynamics (Fig.3C and D).

### Phosphorylation pattern, and not just net-charge of the PLD, influences intracluster dynamics

Our results so far demonstrate the efficacy of phosphorylation as a tunable handle for intra-cluster dynamics, with higher modification fractions resulting in liquid-like clusters. However, even for the same modification fraction, P_frac_, we observe variability in inter-chain contacts and diffusion constants (Fig 2 and 3) for different phosphorylation patterns. To further understand how the positioning of phosphorylated phosphosites could influence the behavior of the condensates, we performed simulations with 40-different phosphorylation patterns, each with P_frac_ = 0.5. Strikingly, for the 40 randomly generated phosphorylation patterns, we observe a significant variation in the total inter-peptide contacts within the cluster. As evident from Fig. 4A, we observe a 2-fold variability in inter-molecular contacts between different phosphorylation patterns despite the net charge of the domain due to the modification being the same. This variability in inter-molecular contacts also gives rise to a 2-fold variation in diffusion coefficients of PLD peptides within the clusters (Fig.4B). This broad distribution in contact densities indicates that the interaction network and intra-cluster dynamics depends not just on the net negative charge of the domain but also on the location of phosphosites.This pattern-dependence of intra-cluster dynamics gives rise to an interesting evolutionary question. Are phosphosite locations non-random and correlated with the location of amyloid hotspots in the FUS prion-like domain? To address this question, we first plot the amyloidigenic propensity of FUS using the ZipperDB algorithm^49^ which computes the compatibility of hexapeptide stretches within protein sequences with the amyloid cross-beta architecture. Interestingly, the amyloidigenicity profile for the unmodified human FUS PLD sequences shows 8 stretches (labelled C1-C8) with ZipperDB energy lower than the reference template, the GNQQNY peptide (Fig.5A). The fully phosphorylated sequence (Fig.5B), with S/T → E shows a complete abrogation of amyloid propensity of the PLD.

**Figure 4:**
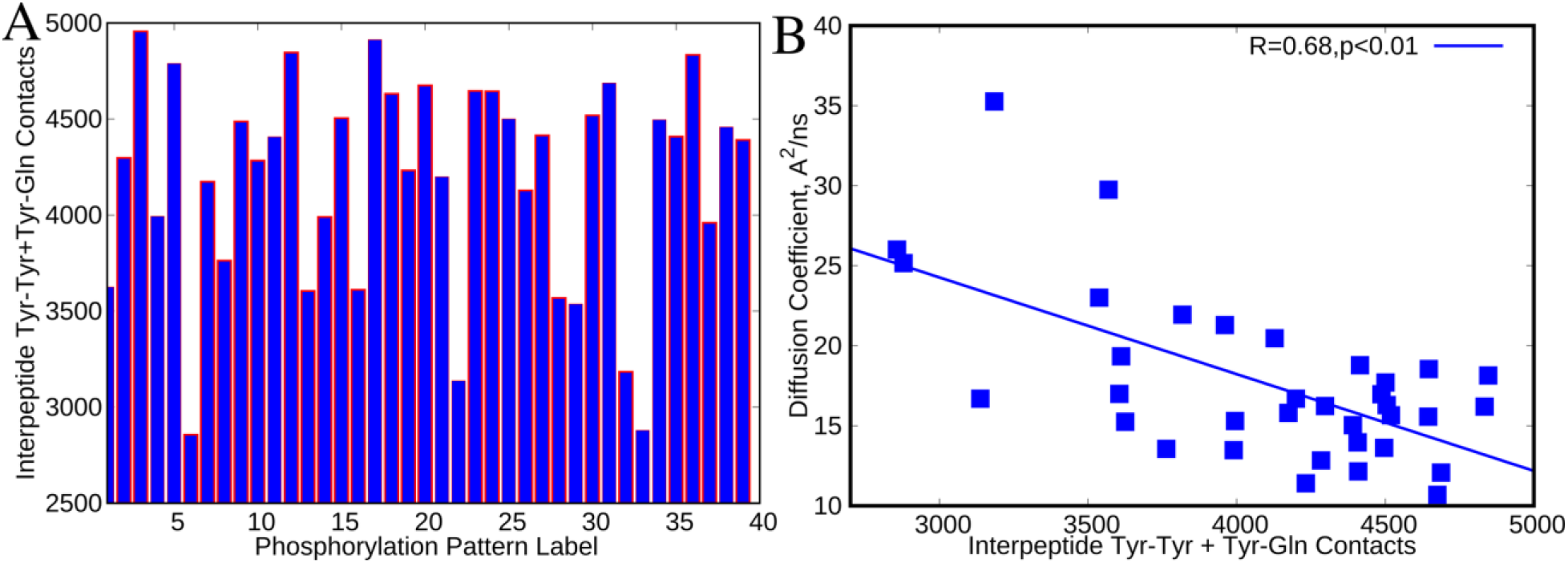
Phosphorylation patterns influence inter-peptide contact probability. A) Inter-chain contacts involving Tyrosine and Glutamine residues, for PLD sequences with different modification patterns with 50% of phosphosites modified (P_frac_ = 0.5). B) Intracluster diffusion coefficients show a negative correlation with inter-peptide contacts involving Tyr and Gln. We observe a 4-fold variability in intra-cluster dynamics, for different modification patterns

**Figure 5:**
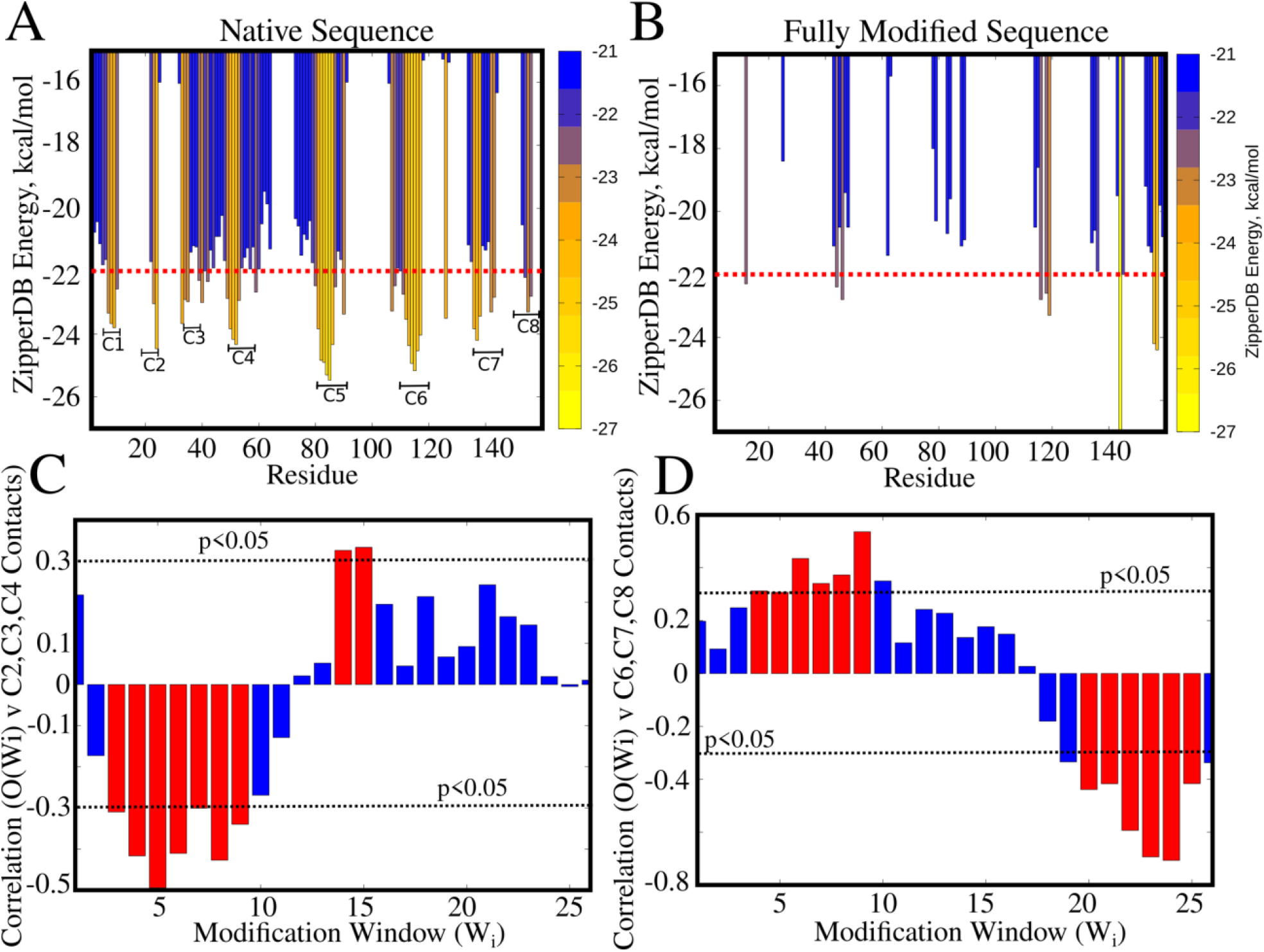
Phosphorylation sites reduce likelihood of amyloidigenic contacts locally. Amyloidigenicity profiles for A) unmodified human FUS PLD, and B) fully phosphorylated FUS PLD sequences. The ZipperDB energy (kcal/mol) indicates the compatibility of a hexapeptide stretch within the protein to the steric-zipper amyloid architecture. Values lower than −22 kcal/mol indicate that the hexapeptide stretch is compatible with the cross-beta structure. C) and D) Phosphorylation patterns are described by 26 overlapping modification windows of 5 consecutive phosphosites -- W_i_. For e.g, the modification window W_1_ comprises of phosphosites at S3,T7,T11,T19,S26 whereas the second modification window W_2_ comprises of phosphosites T7,T11,T19,S26,S30. The occupancy of any modification window O(W_1_) to O(W_25_), for any phosphorylation pattern, is the fraction of residues in the window that are modified. O(W_i_) can, therefore, take values between 0 and 1. Every phosphorylation pattern can thus be defined as a string of occupancy values [O(W_1_), O(W_2_)…..O(W_25_)]. The extent of correlation between different occupancies (O(W_1_) to O(W_25_)) and the corresponding to the sum of amyloid core contacts involving the C1,C2,C3,C4 regions is shown in (C) and C6,C7,C8 in (D). The dotted line respresents the point beyond which the correlations are statistically significant.

We further investigate why different phosphorylation patterns result in a variability in inter-chain contacts and diffusion coefficients, despite the sequences featuring the same net charge. In other words, we test the following hypothesis – if the effect of phosphorylation were merely net-charge driven, any phosphorylation pattern would result in a reduction of contacts involving all 8 amyloid cores (C1 to C8 in Fig.5A) to a similar extent. To address this question, we define a quantity – occupancy of modification window --O(Wi) – that describes different phosphorylation patterns. Here, phosphorylation patterns are described by 26 overlapping modification windows of 5 consecutive phosphosites. For instance, the modification window W_1_ comprises of phosphosites at S3,T7,T11,T19,S26 whereas the second modification window W_2_ comprises of phosphosites T7,T11,T19,S26,S30. Similarly, the W_25_ is defined by phosphosites S129,S131,S135,S142,S148. The occupancy of any modification window O(W_1_) to O(W_25_), for any phosphorylation pattern, is the fraction of residues in the window that are modified. O(W_i_) can, therefore, take values between 0 and 1. Every phosphorylation pattern can thus be defined as a string of occupancy values [O(W_1_), O(W_2_)…..O(W_25_)]. We first compute the correlation between the occupancy of any given window (for 40 different sequences with Pfrac = 0.5) and the amyloid core contacts involving different amyloid cores C1-C8. For instance, in Supplementary Fig.S6A, we show a scatter plot for occupancy of W25 (S129,S131,S135,S142,S148) versus amyloid core contacts involving C6, C7, C8 (region 115-150 in Fig.5A). As evident from Fig. S6A, we observe a statistically significant negative correlation between occupancies in this modification window and the amyloid core contacts involving C6, C7 and C8 cores. On the other hand, the occupancy of W1 shows no such correlation (Fig. S6B). In Fig.5C and D, we plot the extent of correlation between different occupancies (O(W_1_) to O(W_25_)) and the sum of corresponding amyloid core contacts involving C1,C2,C3,C4 (Fig.5C) and C6,C7,C8 (Fig.5D).

Interestingly, an increased occupancy of modification windows W_1_ to W_10_ results in a statistically significant negative correlation with amyloid core contacts involving C2,C3 and C4. For windows labeled W11 to W24 we either observe a weak positive or statistically insignificant correlation for C2,C3,C4 contacts. In other words, to reduce the density of contacts involving C2,C3,C4, the modification patterns must enrich phosphosites that are localized in the N-terminal region of the protein. Similarly, to reduce the likelihood of amyloid core contacts involving the C6, C7 and C8 regions, the phosphorylation patterns must involve residues that comprise modification windows W_20_ to W_25_, i.e the region between residues 110-140 of the protein. In fact, for patterns which have low occupancies in W20 to W25 and higher occupancies for W4 to W10, we observe an increase in C6, C7, C8 contacts, as seen from a statistically significant positive correlation for these regions in Fig.5D. Our results suggest that an effective phosphorylation pattern must, therefore, include modification sites that are in the vicinity of each potential amyloid core (C1 to C8) in the FUS_1-165_ sequence.Our coarse-grained simulations reveal a strongly local, position specific effect of phosphorylation.

While the coarse-grained simulations can capture the effect of phosphomimetic substitutions on inter-peptide contacts, the coarse-grained model cannot capture secondary structural transitions. We therefore performed all-atom molecular dynamics simulations with different amyloid-core segments from FUS_1-165_ and studied the difference in secondary structural propensity of various amyloid core peptides (Supplementary Table S4) – i) 3-13 (Core C1), ii) 47-57 (Core C4) iii) 79-95 (Core C5) iv) 105-120 (Core C6) v) 135-141 (Core C7) – in the presence and absence of phosphomimetic substitutions. In order to study the local effect of phosphorylation on the amyloid core contacts, we introduce a negative charge via a Glutamic-acid substitution at the phosphosite location (blue bars with labels tagged P in Figure 6). Using metadynamics simulations we drive the self-assembly of peptides into large clusters at simulation timescale (100 ns). We then analyze the secondary structural content of the system by computing the proportion of all residues in the system in the beta-sheet region of the Ramachandran plot (across 50 copies of the peptide in the simulation box). Strikingly, while the β-sheet content for the wild-type peptides can be as high as 50%, introduction of phosphomimetic substitutions results in a reduction of β-sheet content by more than half, in all the peptides under study. The β-sheet content for phosphomimic sequences (tagged P) approaches that of the polyG sequence which has natively low propensity for β-sheets. We further analyzed the secondary structural propensity using the Ramachandran number^50^ – a secondary structural order parameter that collapses the 2-dimensional information from the Ramachandran plot (Eqn. 4). In Fig.6B and Supplementary Fig. S7 we show the Ramachandran number plots for various amyloid core peptides and their phosphomimetic counterparts.

**Fig 6.**
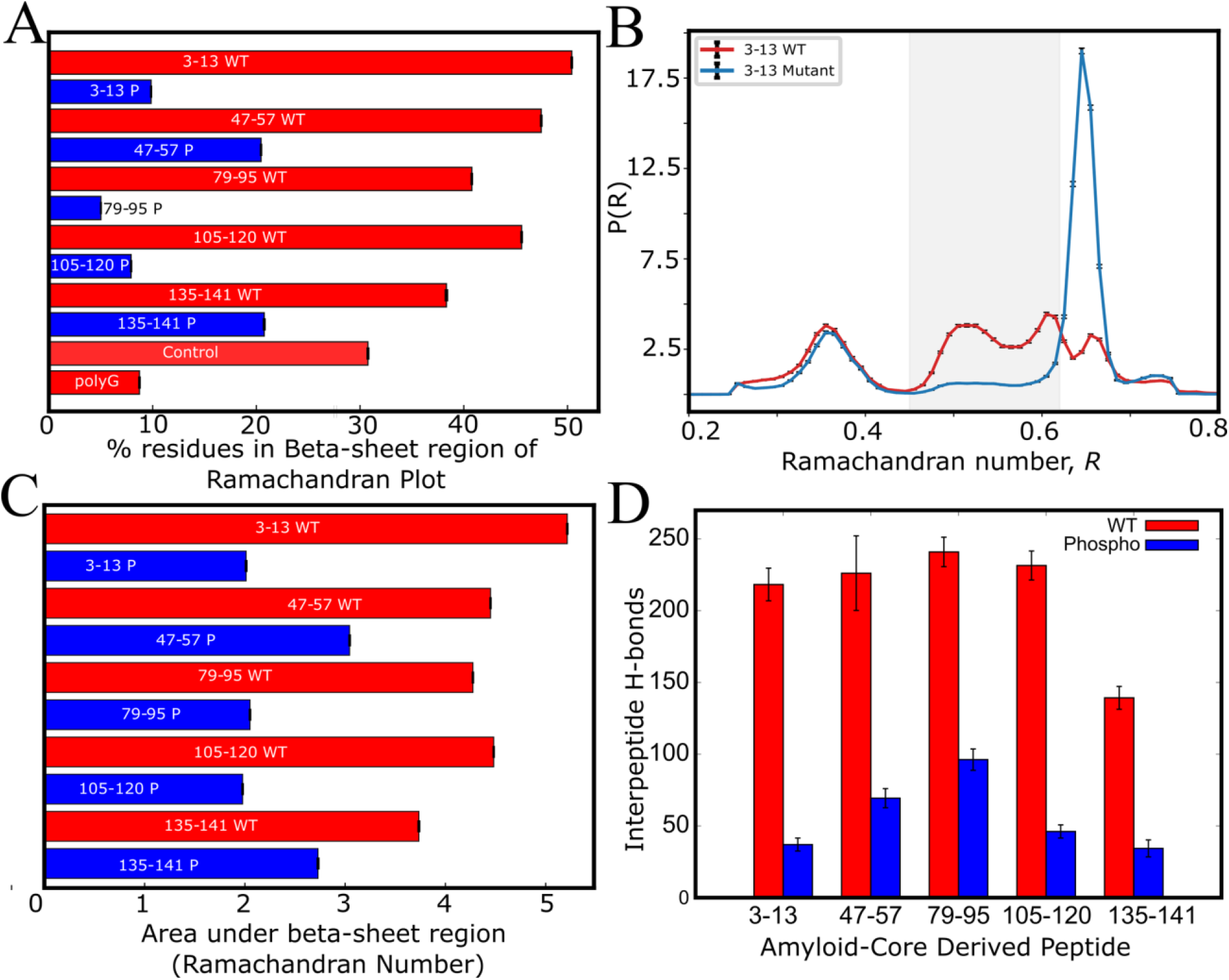
Accessibility of β-sheet region of the Ramachandran plot for amyloid core-derived peptides. A) Proportion of residues in the β-sheet region of the Ramachandran plot for self-assembly simulations with 50 copies of peptides corresponding to different amyloid core regions (C1-C8 in Fig.5). B) Area under the curve in the Ramachandran number plots. The Ramachandran number, R(φ,ψ) = (φ+ψ+2π)/4π. The red bars correspond to simulations with native peptide sequence (WT) while the blue bars correspond to phosphomimetic variants of the segment, with phosphosite residues of the WT modified to Glutamic acid. Introduction of a negative charge at the phosphosite location via glutamic acid substitution results in a significant decrease in the accessibility to the β-sheet region of the Ramachandran plot. D) Mean inter-peptide hydrogen bonds for the amyloid core-derived peptides. Wild type peptides are shown in red while the phosphorylated variants are shown in blue.

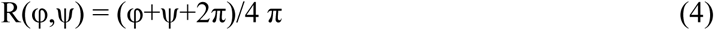

The Ramachandran number for any residue in equation 4 ranges from 0 to 1, with the combination of φ,ψ angles corresponding to the β-sheet region resulting in R(φ,ψ) values between 0.45 to 0.65 (shaded region in Fig. 6B,C). As evident from the Ramachandran number plot in Supplementary Fig. S6, the area under the β-sheet region of the Ramachandran number curve goes down for phosphomimic substitutions, for all peptides under study (Fig. 6B,C and Supplementary Fig.S7). These results establish that introduction of a negatively charged phosphomimetic substitution in the vicinity of the amyloidogenic region results in a reduced accesibility to the β-sheet of the Ramachandran plot. Further, the phosphomimetic variants also display a dramatic reduction in the ability to form interpeptide hydrogen bonds (Fig. 6D and Supplementary Fig. S8). This is consistent with the observation from metadynamics simulations that show that introduction of phosphomimetic substitutions destabilizes the self-assembled state altogether (Fig. S9). These results indicate that phosphorylation can be an effective local checkpoints against inter-peptide β-sheet formation, a signature of the amyloid-like state.

The Ramachandran plot analysis for the amyloid-core peptides that were studied using atomistic simulations are only used as a measure of the accessibility to the beta-sheet region of the Ramachandran plot. Being able to access this region of the Ramachandran plot is the first pre-requisite to formation of beta-sheets. In this context, the more striking and relevant result is that phosphomimic mutations to these peptides results in an abrogation of accessibility to the beta-sheet region of the R-plot. Therefore, while the results in Fig.6 are not sufficient to claim that the WT peptides assemble into amyloid-like beta sheets in our simulations, they show a dramatic reduction in the accessibility to the beta-sheet region along with drastic reduction in interpeptide hydrogen bond formation (Fig.S8), necessary conditions for amyloid-formation.

### Evolution selects phosphosites in the vicinity of FUS amyloidigenic regions

PLD sequences with the same modification fraction but different phosphosites can exhibit variability in inter-protein contacts as well as intra-cluster dynamics (Figs. 3-5). The local effect of phosphorylation, as opposed to a net-charge effect gives rise to a crucial question. Could phosphosite loci, have evolved to reduce the overall amyloidigenic propensity (across different amyloid cores) in the PLD of FUS proteins?

To test this hypothesis, we investigate the amyloidigenic propensity and the corresponding phosphosites in 85 different mammalian FUS PLD sequences (Fig 7A). Interestingly, we observe a strong correlation between the native amyloidigenic propensity and the potential number of phosphosites in a mammalian FUS PLD sequence. This result suggests that PLD sequences that are inherently more prone to solidification via amyloid-like interactions more likely require phosphosites to prevent possible aberrant transitions. We further simulate these 85-mammalian FUS PLD sequences using the coarse grained model and observe a statistically significant correlation between sticky interactions (Y-Y and Q-Q) within the cluster and the potential phosphosites in the sequence (Fig.7B). Importantly, different mammalian PLD sequences also show dramatically different intra-cluster dynamics, with the native primate PLD sequences which harbor the most number of phosphosites showing the slowest intracluster diffusion (Fig.7C). PLDs of mammalian sequences with fewer number of phosphosites were more likely to assemble into liquid-like clusters. Consistent with this hypothesis, sequences which are more prone to forming Q-Q and Y-Y contacts also exhibit slower intra-cluster dynamics in our simulations (Fig. 7D).

**Figure 7:**
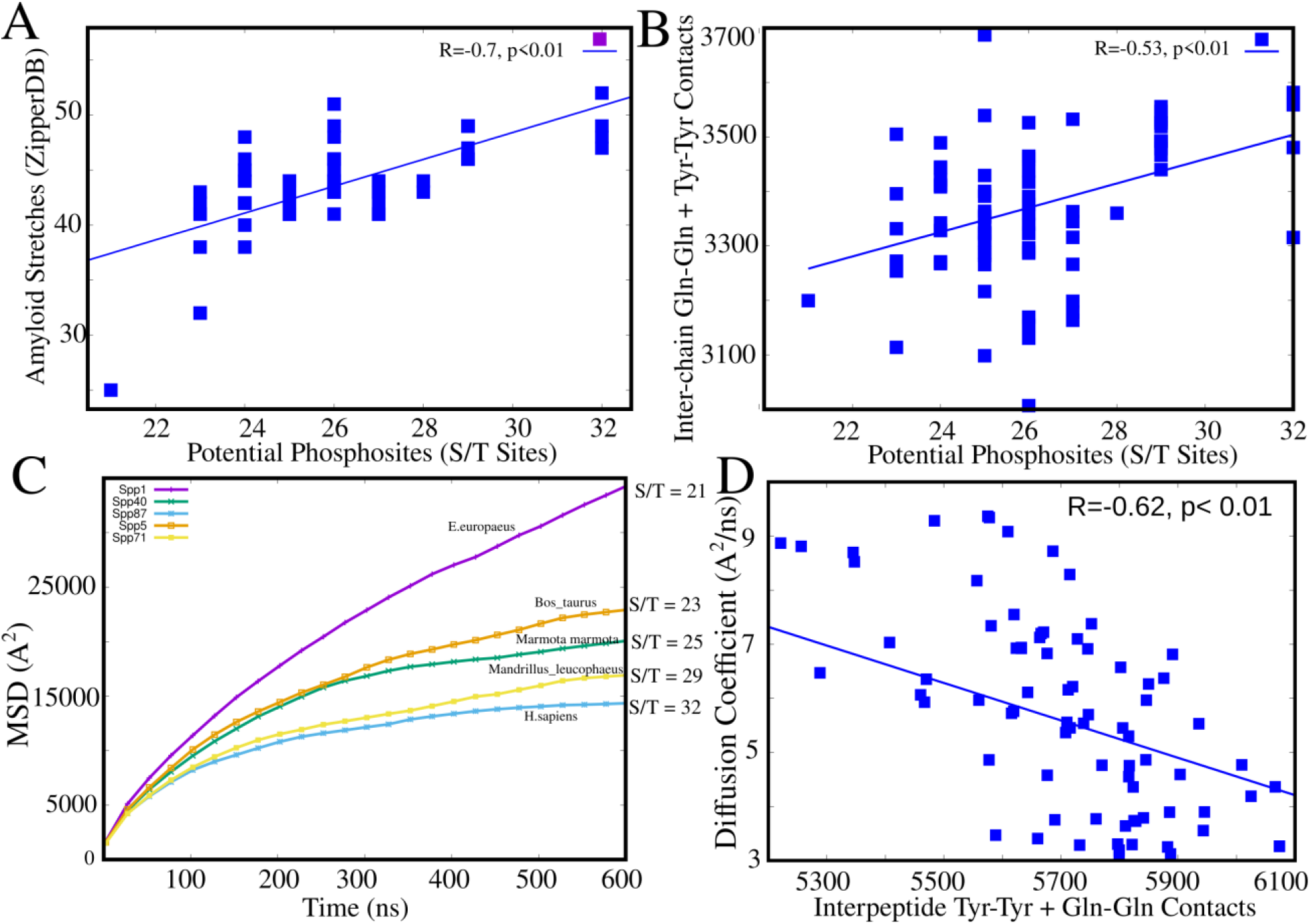
Evolutionary enrichment of phosphosites to ensure liquidity of condensates. A) Correlation between the total number of cross-beta amyloid-compatible stretches in a mammalian FUS PLD sequence and the total number of potential phosphosites in the same sequence. A statistically significant correlation of 0.7 was observed (p<0.01). B) Correlation between the number of phosphosites in any mammalian sequence, and the total number of sticky interactions (involving Tyr and Gln) within the self assembled cluster (computed from simulations). Each datapoint corresponds to a different mammalian FUS PLD sequence. C) Mean square displacement profiles for different representative mammalians sequences with different number of potential phosphosites. D) Mammalian sequences that are more prone to intra-cluster Q-Q,Y-Y contacts show less dynamic behavior in simulations.

Since amyloid-forming regions increase the propensity of protein-protein interactions in PLD sequences, phosphorylation sites may serve as checkpoints against liquid-solid transition via amyloid-prone regions in low complexity domains of these proteins. Since amyloid-forming regions increase the propensity of protein-protein interactions in PLD sequences, phosphorylation sites may serve as checkpoints against liquid-solid transition via amyloid-prone regions in low complexity domains of these proteins. To test this hypothesis, we compared the number of amyloid-prone regions in the human FUS PLD domain with a neutrally-evolved protein sequence of 4000 amino acids. Briefly, we simulated neutral protein sequence evolution using the codon composition that matches human FUS PLD sequence, and employed realistic divergence times of FUS in mammals (Figure 8A, see Methods for details). We then calculated the distance of phosphosites to the amyloid-forming residues (ZipperDB energy < −22 kcal/mol) in the neutrally-evolved FUS sequence and compared them with those found in the human FUS PLD sequence. As shown in Figure 8B, the distance between phosphosites and amyloid-forming regions was significantly shorter in human FUS PLD sequence compared to the neutrally-evolved FUS protein (*p*=0.0056; One-sided Wilcoxon rak sum test).

**Figure 8.**
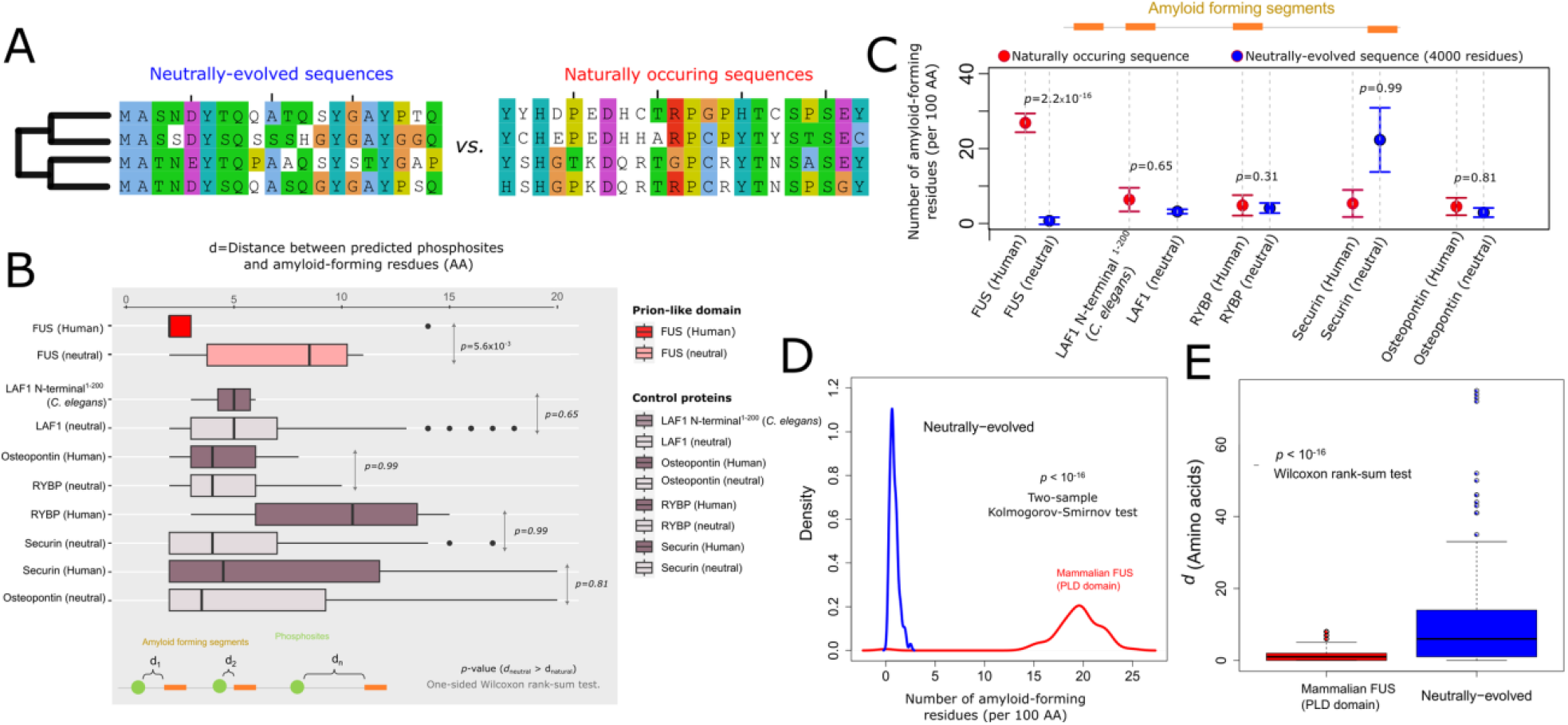
The number of amyloid-forming regions and their proximity to potential phosphorylation sites have evolved under positive selection. A) We compared the number of amyloid forming regions and the distance of phosphosites to such regions in naturally occurring protein sequences, and sequences that had evolved under null hypothesis of neutral evolution (see Methods for details of simulations). B) The distance between potential phosphosites and predicted amyloid forming residues in the FUS PLD and several control proteins. We selected *C. elegans* LAF1 as an example of a phase-separating and non-aggregating protein and three fully disordered and non-phase separating human proteins (RYBP, Securin, and Osteopontin). C) The number of amyloid forming residues (ZipperDB energy < −22 kcal/mol) per 100 amino acids in the PLD domain of Human FUS and in the N-terminal disordered domain of LAF1 (residues 1-200), and in the rest of control proteins. For these comparisons (panels B and C), for each protein we used a single neutrally evolved sequence with the length of 4000 amino acids as comparison with null hypothesis of neutral evolution (see Methods). The p-values on panels B and C are calculated from one-sided Wilcoxon rank sum test with the alternative hypothesis that the distances between phosphosites and amyloid forming regions are shorter in naturally-occurring sequences (panel B), and neutrally-evolved sequences have fewer amyloid-forming residues per 100 amino acids (panel C). D) The density of the number of amyloid forming residues (ZipperDB energy < −22 kcal/mol) in 85 mammalian FUS PLD sequences (shown in blue), and neutrally evolved sequences (shown in red). E) The distance between potential phosphosites and amyloid forming residues in mammalian FUS PLD sequences (shown in blue), and neutrally evolved sequences (shown in red). For comparisons in panels D and E, we used 100 neutrally evolved sequences each with the length of 600 amino acids. We predicted the phosphorylation sites using NetPhos v3.0 server.

We further compared the distribution of distances between putative phosphosites and amyloid-forming regions in naturally occurring sequences of several control proteins and their corresponding neutrally evolved sequences (see Methods). Our first control protein was the phase-separating and non-aggregation-prone protein LAF-1. This protein is an ATP-dependent RNA helicase and consists of helicase and disordered domains. We selected the N-terminal disordered region of LAF-1 (residues 1-200) that is necessary and sufficient for the phase-separation of this protein^51^. We also selected disordered and non-phase separating proteins as further controls in our comparison. Specifically, we selected three intrinsically disordered proteins: Osteopontin, Securin, and RYBP. Osteopontin is one of the major components of mineralized extracellular matrices of bones^52^. Securin is involved in the control of metaphase-anaphase onset, and RYBP is part of the Polycomb group multiprotein PRC1-like complex that represses the transcription of many genes during development. Importantly, these proteins require phosphorylation for their physiological function making them suitable candidates for our analyses^53,54^. Like the case of FUS PLD, we identified the amyloid forming regions in these proteins using the ZipperDB algorithm and considered any residue whose ZipperDB energy <-22 kcal/mol as an amyloid-forming residue. We predicted phosphorylation sites using the NetPhos 3.0 webserver. To exclude the putative S/T residues that form the core of amyloid fibrils in these proteins, we only considered phosphosites that were more than one amino acid away from amyloid-forming residues. Interestingly, the distances between putative phosphosites and amyloid-forming regions were only significantly shorter in the case of FUS prion-like domain compared to our control proteins (Figure 8B; one-sided Wilcoxon rank sum test). Likewise, the number of amyloid-forming residues per 100 amino acids was substantially higher only for the prion-like domain of FUS compared to control proteins (*p*=2.2×10^−16^, one-sided Wilcoxon rank sum test; Figure 8C). This difference was insignificant for LAF-1, RYBP, Securin, and Osteopontin with p-values ∼ 0.65, 0.31, 0.99, and 0.81, respectively (one-sided Wilcoxon rank sum test). These observations show that both the number of amyloid-forming residues and their proximity to phosphosites have evolved under positive selection in human FUS, compared to our control disordered proteins. Having established special role of phosporylation to control phase state of human FUS, we extended the evolutionary analysis of the PLD domain of FUS to 85 mammalian FUS sequences to see whether our observation of positive selection in the number of amyloid-forming residues and their proximity to phosphosites can be extended to other mammalian FUS sequences. As before, our null hypothesis for this analysis was that phosphosites and amyloidegentic sites evolve neutrally on their corresponding phylogenetic trees. To that end, we generated 100 neutrally-evolved sequences of 600 amino acids each as described in Methods. Indeed, the number of amyloid-forming residues per 100 amino acids was ∼ 19.38 ± 2.91 in mammalian sequences which was significantly higher than the number of amyloid-forming residues in control neutrally-evolved sequences (=3.15 ± 1.62; *p* < 10^−16^; Two-sided Kolmogorov-Smirnov test, Figure 8D). The distance of such residues to phosphosites was also significantly shorter in mammalian PLD sequences compared to neutrally evolved sequences (Figure 8E, *p* < 10^−16^; Wilcoxon rank-sum test). Indeed, an order of magnitude increase in the number of neutrally evolved residues in the simulations (100×600=60000 AAs) compared to 4000 AAs in Figure B substantially increased the significance of our observation of shorter distances between phosphosites and amyloid-forming residues in mammalian FUS proteins, compared to neutrally evolved sequences. We also looked at the local sequence environment close to the amyloid-forming regions in mammalian PLD sequences and found that such sequences were significantly enriched in dipeptides containing serine compared to neutrally evolved sequences making them suitable kinase motifs (Supplementary Figure S10). Altogether, these observations show that evolution has selected phosphorylation sites near amyloid-forming regions in FUS likely as checkpoints to minimize amyloid formation and liquid-to-solid phase transition.

## Discussion

While being efficient promoters of phase-separation, IDRs are also often extremely prone to aberrant, irreversible aggregation into amyloid-like structures.^55^ The ability of proteins harboring PLDs to switch between different interaction modes could help them access diverse states including the liquid-like and amyloid-like configurations. Living systems, therefore, operate at the edges of biomolecular phase-transitions, with small perturbations often resulting in a dramatically altered state of a system. Studying the mechanisms that allow cells to exploit the inherent self-assembling propensity of IDRs while guarding against aberrant phase transitions can help shed light on biomolecular function and cellular evolution.

The FUS protein is a well studied phase separating protein that has been observed to assemble into dense compartments in response to DNA damage. FUS harbors long stretches of intrinsically disordered regions, and is known to undergo liquid-solid transitions, a process that is accelerated in presence of ALS mutations. These factors make FUS a useful model protein to study proteostatic mechanisms accompanying functional protein-self assembly. Rhoads et al reported that FUS is multiphosphorylated in response to DNA damage, with 28 putative phosphosites showing signatures of phosphorylation under different conditions^17,18^. Crucially, not all phosphosites get modified at the same frequency, resulting in differential phosphorylation of FUS under varying conditions. This gives rise to an interesting biological implication – could phosphorylation patterns in partially phosphorylated FUS be exploited by cells for granular control over droplet dynamics? Does evolution select for phosphosites at specific locations along the FUS PLD sequence to prevent liquid-solid transitions? Using coarse-grained simulations of over 100 different phosphorylated sequences which vary in extent as well as pattern of phosphorylation, we establish that the effect of phosphorylation is not merely net-charge driven. While sequences that are extensively phosphorylated (> 60% phosphosites modified) show complete abrogation of self-assembly, partially modified FUS (<60% phosphosites modified) remains stable in the dense phase. Crucially, even in this partially phosphorylated regime, we observe a 4-fold variation in diffusion coefficients within the dense phase depending on the extent, and pattern of phosphorylation (Figure 3C and 4B). Our simulations suggest that the altered intradroplet dynamics for sequences with the same extent of phosphorylation but different pattern is an outcome of variability in sticky interpeptide contacts involving Y/Q residues in the dense phase. Overall, our simulations show that the effect of phosphorylation goes beyond an alteration of the net charge of the PLD. Correlations between presence of modifications at different locations and the net amyloid-prone contacts suggest that the effect of phosphorylation is highly local (Fig. 5). Given the presence of several amyloid-prone regions in the FUS PLD, an overall reduction in amyloid-propensity would necessitate patterns which include phosphorylation across different modification windows (Fig. 5).

The results from coarse-grained simulations give rise to an interesting mechanistic hypothesis. To efficiently abrogate amyloidigenicity while maintaining condensed phase, evolution must place phosphosites to spatially correlate with the location of amyloid-prone regions in the PLD sequence.

In order to test this hypothesis, we studied 85 mammalian sequences for their amyloid propensity profiles as well as inter-peptide contacts using Langevin dynamics simulations. Dasmeh et al^45^ have previously established that the PLD of FUS is the region of the protein that shows the highest degree of sequence entropy in the alignment, with *S* → *G* and *G* → *S* being the most abundant substitutions in the 85 mammalian PLD sequences. *G* → *S* transitions resulting in phosphosites is 3-fold more likely in primate FUS PLD sequence as compared to their other mammalian counterparts. Primate sequences which harbor the greatest number of phosphosites also result in clusters with a greater number of Y/Q contacts (Fig. 6B). In general, the mammalian FUS sequences that result in less dynamic assemblies were also more likely to contain greater number of potential phosphosites (Fig.6A, C, D). Positive selection for phosphosites in primate FUS PLD sequences could therefore be an evolutionary mechanism to ensure the dynamicity of structures prone to amyloid formation (Fig.6C).

Overall the results of molecular simulations, are suggestive that phosphorylation patterns should be non-random to serve as effective checkpoints against amyloid formation in FUS. Therefore, we infer that amyloid-prone sequences get utilized by cells to promote protein clustering at lower threshold concentrations while the potential drawback of this mechanism - aberrant transitions to irreversible solid-like structures - are prevented by phosphorylating residues in the vicinity of these amyloid cores.Therefore in natural sequences such patterns might have evolved under the selection pressure to minimize several amyloid-prone contacts. To test this hypothesis we turned to a bioinformatics study where we compared statistics of amyloid and phosphorylation sites in natural FUS proteins with model sequences generated under the null model hypothesis of neutral evolution. Our evolutionary analysis showed that mammalian FUS sequences are indeed significantly more likely to harbor amyloidigenic sites compared to neutrally evolved sequences with same codon frequencies as mammalian FUS sequences (Fig.8 and Supplementary Fig.S7). Further, the distance between amyloidigenic sites and the nearest phosphosite was also significantly shorter in mammalian FUS sequences, as compared to the neutrally evolved counterparts with no selection pressure. Significantly, such an enrichment of amyloidigenic sites and the accompanying phosphosites was specific to FUS and not observed in case of three fully disordered and non-phase separating human proteins (RYBP, Securin, and Osteopontin) as well as a phase-separating protein LAF1 which does not undergo liquid-solid transitions (Fig 8B and C). These results are in conceptual agreement, with the findings by Kumar et al which show that PTMs have negligible effect on abrogation of fibrilation if they are not in close proximity to the amyloid core^16^.

Evolution, therefore, uses phosphorylaton as a switch that could help cells toggle the state of the protein-rich cluster, by shifting phase boundaries as well as intra-cluster dynamics of proteins (Fig. 9). This could allow cells to exploit aggregation-prone stretches to promote phase-separation and biomolecular condensation while safeguarding against the detrimental effects of amyloidosis. In summary, our study highlights an elegant and yet subtle evolutionary balancing act whereby amyloid-prone sequences are actually “selected-for” the efficacy in promoting self-assembly of FUS while also selecting for phosphorylation sites in close proximity (not within) to the amyloid-prone regions as checkpoints to prevent “excesses” of self assembly such as solidification in the protein-rich condensate.

**Figure 9.**
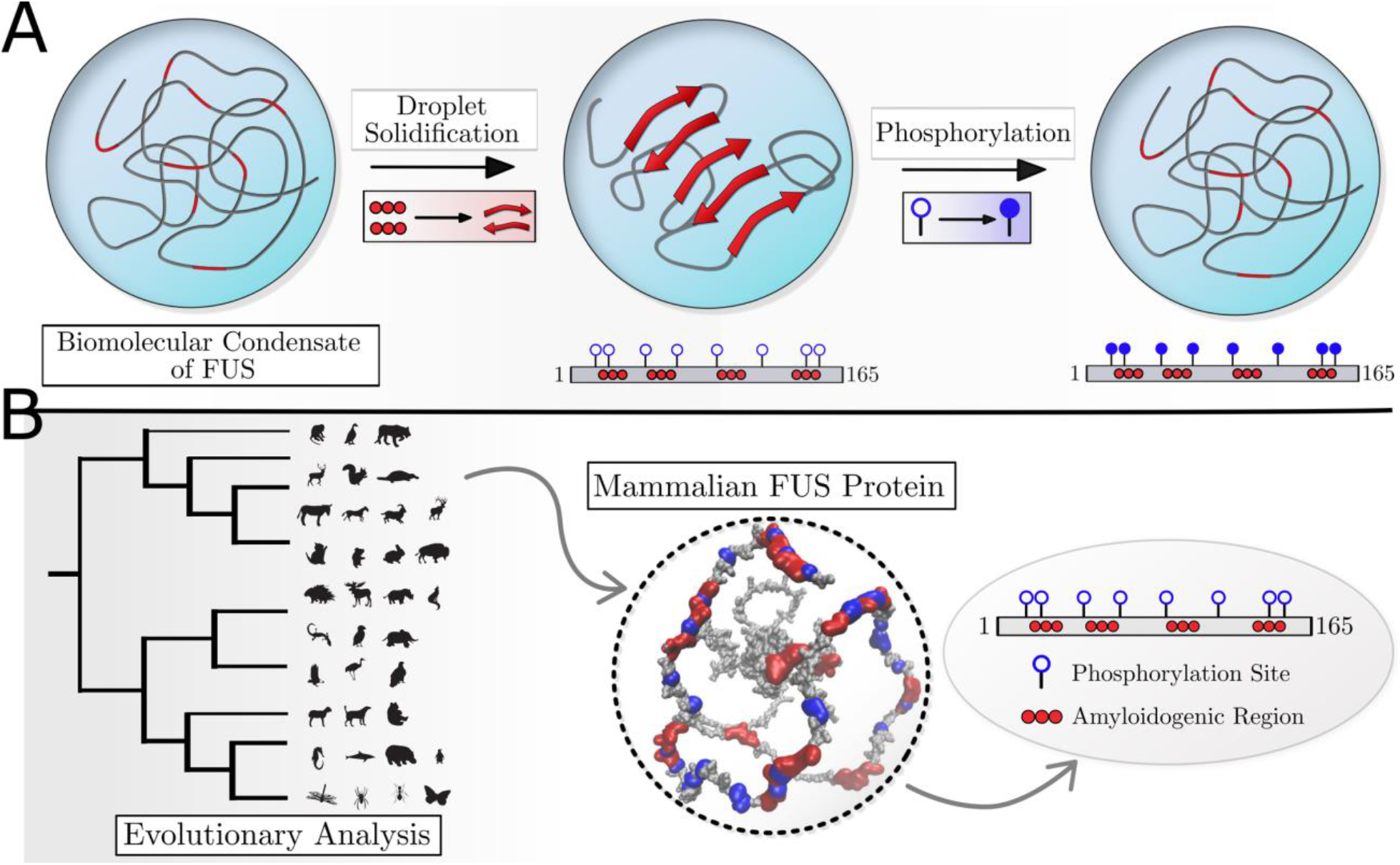
Graphical summary of the study. A) Coarse-grained simulations show that phosphorylation results in a decrease in sticky Tyrosine and Glutamine contacts that result in a more dynamic intra-cluster environment. B) Evolutionary bioinformatics analysis and coarse-grained simulations with 85 mammalian FUS PLD sequences reveals a strong correlation between the number of phosphorylation sites (purple beads in the ball and stick representation) and the amyloidogenic propensity (red beads in the ball and stick representation) of the native sequence. Comparison with neutrally evolved sequences reveals that mammalian sequences localize phosphosites in the vicinity of amyloid prone regions.

## Acknowledgement

The authors would like to thank Junlang Liu, Eugene Serebryany, David Thorn and Amir Bitran for their suggestions and valuable discussions.

## Author Contribution

SR, PD and ES were involved in the conceptualization of the work. SR, PD, SF and ES wrote the paper. SR designed and performed the coarse-grained simulations. PD setup and performed the bioinformatics study. SF performed the atomistic simulations and analyzed the all-atom simulation data.

