## Supporting Information for "Phosphorylation sites are evolutionary checkpoints against liquid-solid transition in protein condensates" (from another version of this preprint: version 3, 2023-01-13)

### Supplementary Figures

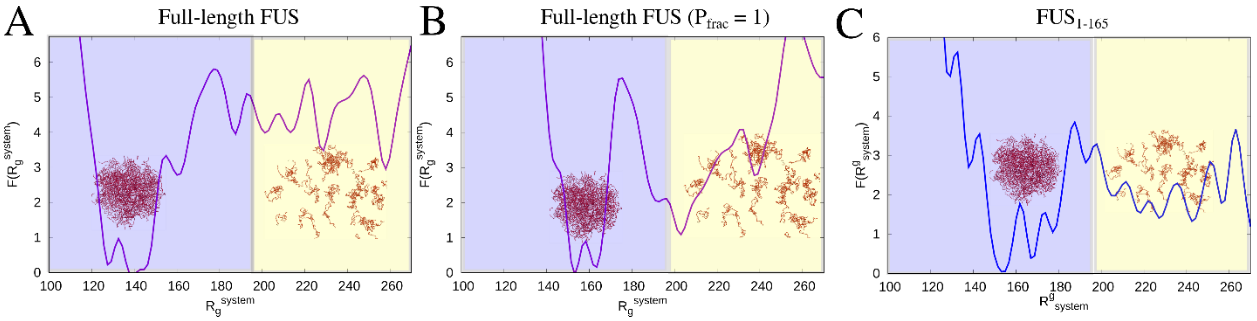


**Figure S1. Metadynamics Free Energy Profile for Full-length FUS, fully-phosphorylated Full-length FUS and FUS$_{1\text{-}165}$ phase separation.** Free energy of a system of 50 FUS chains (effective concentration of 100 uM). The reaction coordinate along which the metadynamics simulation is performed if the radius of gyration of an imaginary polymer that is composed of the 50 center of masses corresponding to the constituent polymer chains in the system. Values of $R^g_{system}$ approaching 150 Angstroms corresponds to a single large spherical condensate while at higher values (> 200 Angstroms), the system approaches the fully mixed state.

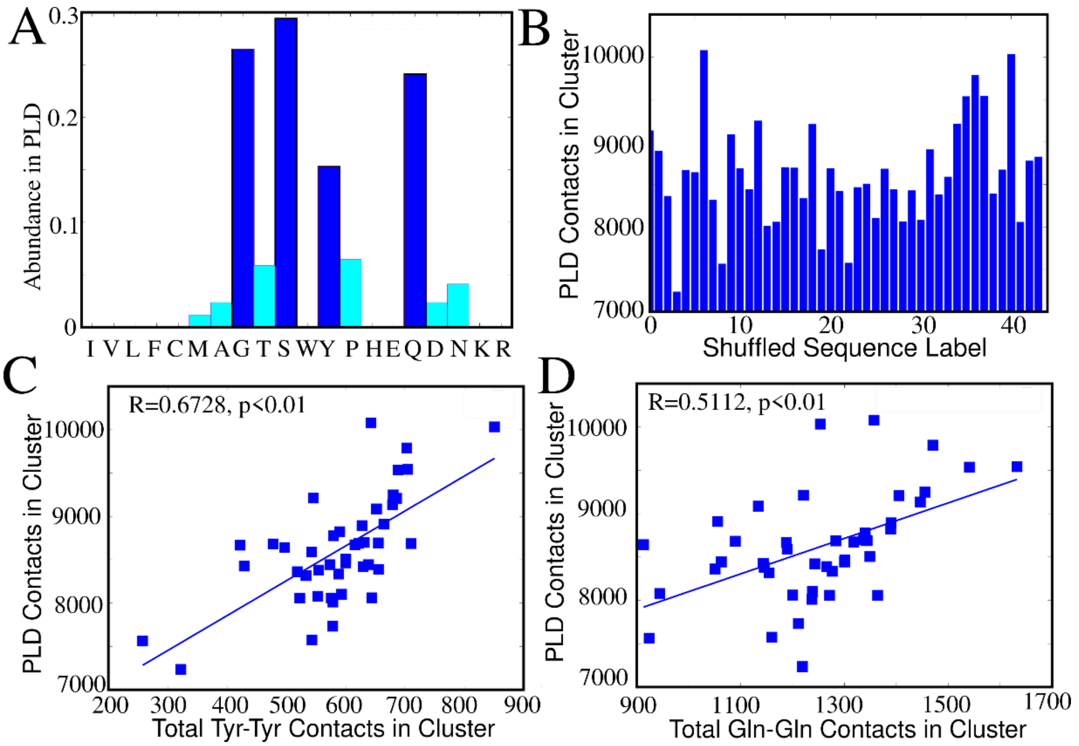


**Figure S2. Identifying Key Contacts.** A) The frequency of occurence of various residues in the human FUS PLD sequence (residues 1-165). B) The mean inter-chain PLD-PLD interactions within a cluster, for a set of 40 FUS sequences with shuffled PLD regions. Each point in the figure corresponds to a different suffled PLD sequence, with RBD remaining the same. C) and D) The total interpeptide PLD-PLD contacts correlate strongly with Tyr-Tyr and Gln-Gln interactions, suggesting their role as sticker residues.

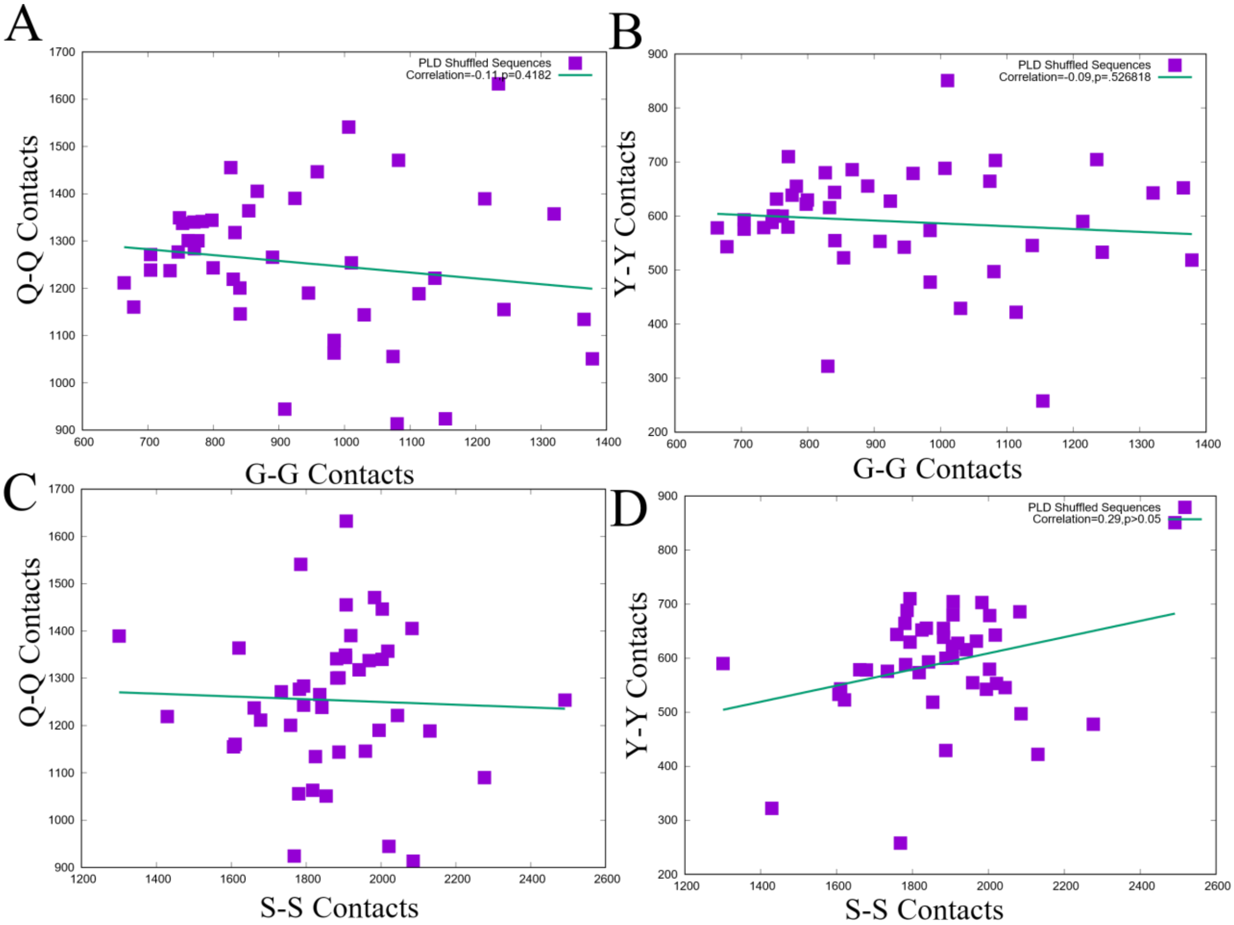


**Figure S3. Identifying Key Contacts.** Increase in Q-Q and Y-Q contacts does not result in an increase in S-S and G-G contacts.

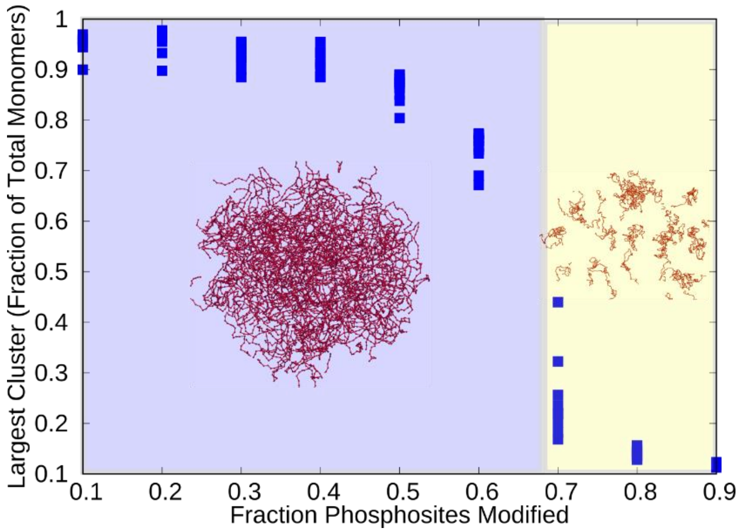


**Figure S4.** Size of largest cluster in the simulation (as a fraction of total monomers) as a function of modification fraction. For modification fraction ($P_{frac}$) of 0.6 or higher, the largest cluster sizes $L_{clus} \rightarrow 0$ suggesting that the system tends to remain predominantly in the monomeric state in this regime.

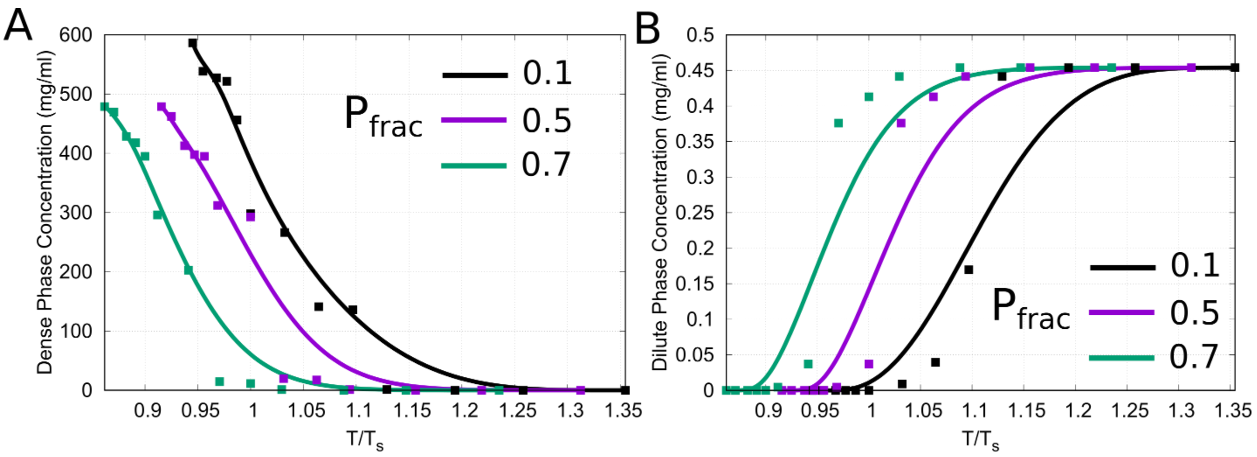


**Figure S5.** Concentration of FUS PLD in A) the dense phase, and B) dilute phase as a function of simulation temperature (T). Ts refers to a temperature of 310K, the standard simulation temperature at which all other simulation results in the paper were reported. The black, purple and green curves represent FUS PLDs with a phosphorylation fraction of 0.1, 0.5 and 0.7, respectively. The solid curves are a guide to the eye.

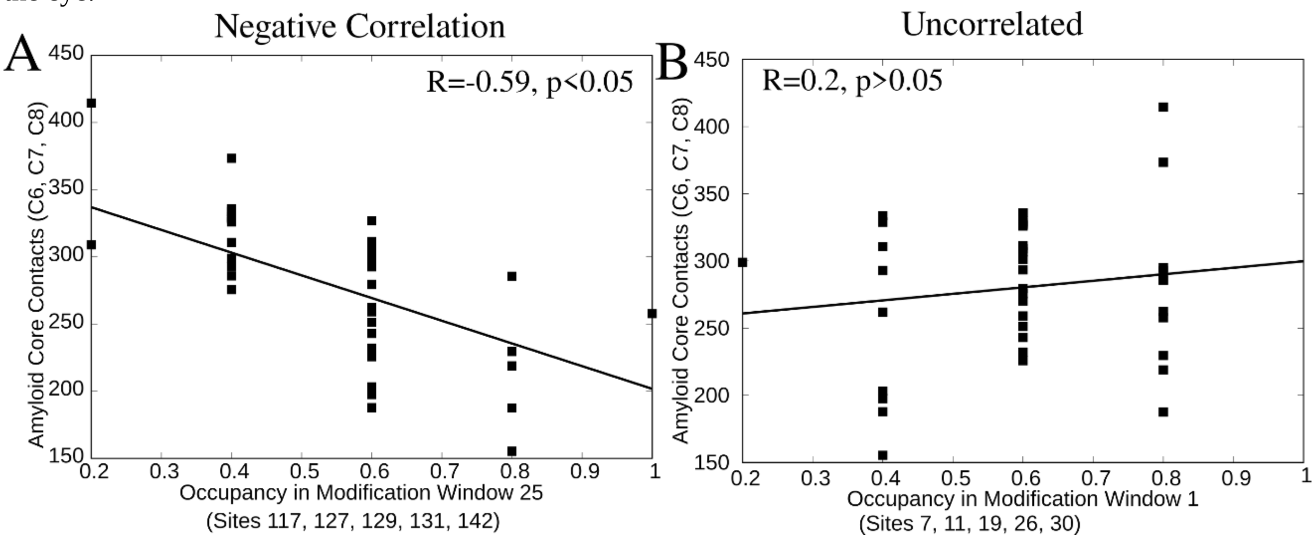


**Figure S6.** A) Scatter plot for occupancy of $W_{25}$ (S117,S127,S129,S131,S142) versus amyloid core contacts involving C6, C7, C8 (region 115-150 of the FUS PLD sequence). We observe a statistically significant negative correlation between occupancies in this modification window $W_{25}$ and the amyloid core contacts involving C6, C7 and C8 cores. B) Occupancy of $W_1$ shows no such correlation with amyloid core contacts involving C6, C7, C8 cores.

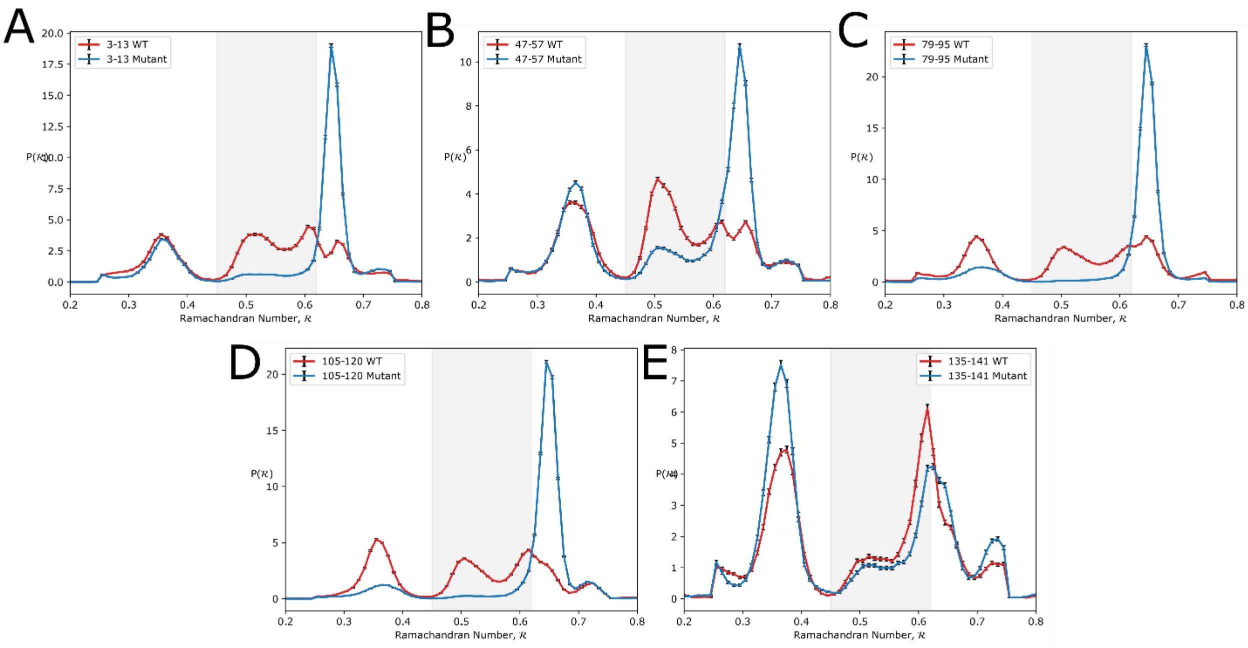


**Figure S7. Ramachandran Number for amyloid-core derived peptides (atomistic simulations).** Frequency of amino acid residues accessing any given Ramachandran number value between 0 and 1**.** The shaded region between 0.45 to 0.62 corresponds to the beta-sheet region of the Ramachandran plot. Panels (A),(B),(C),(D) and (E) correspond to peptides 3-13, 47-57, 79-95, 105-120 and 135-141, respectively.

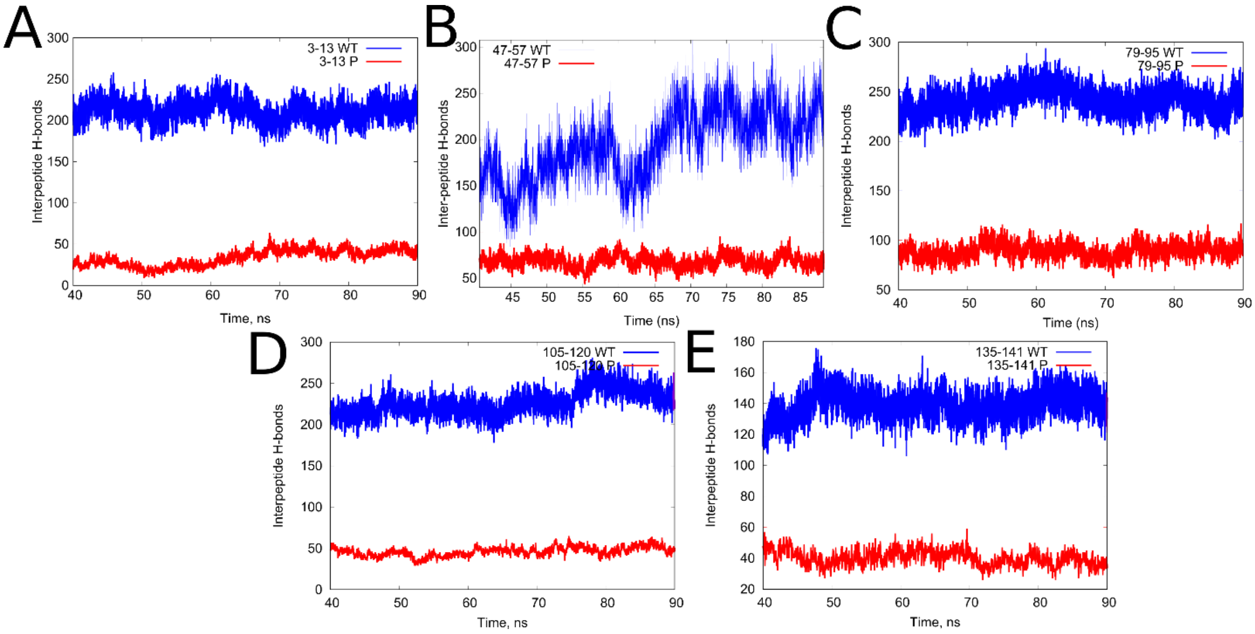


**Figure S8. Interpeptide Hydrogen Bonds for FUS amyloid core-derived peptides.** Panels A), B), C), D) and E) refer to peptides corresponding to regions 3-13, 47-57, 79-95, 105-120 and 135-141, respectively. The blue curves correspond to wild type peptide whereas the red curves correspond to the sequences with phosphomimetic substitutions.

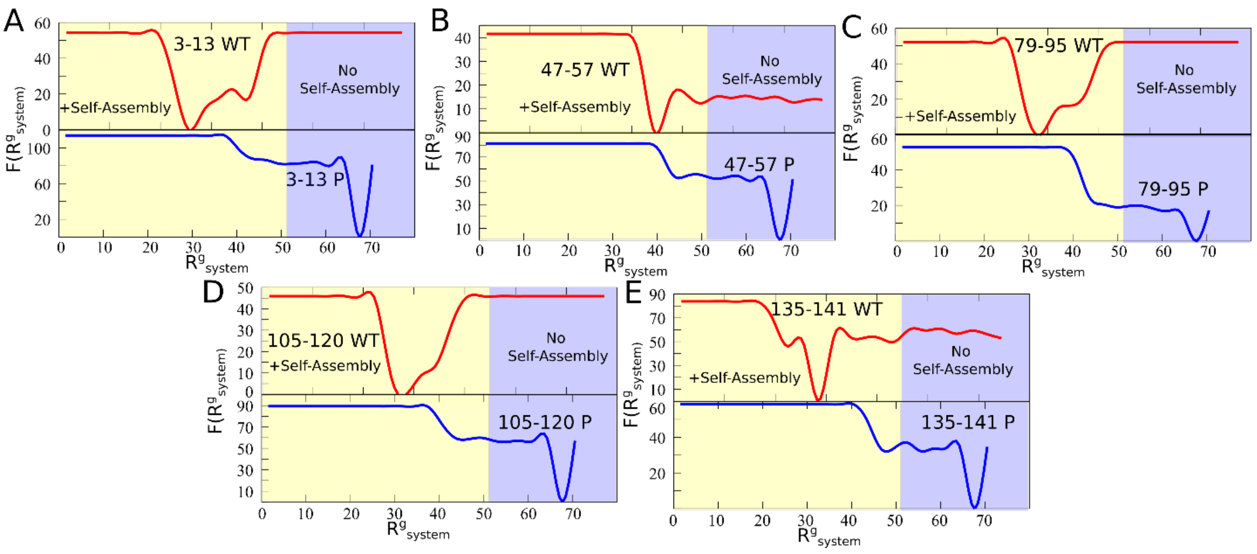


**Figure S9. Free Energy Profiles for Amyloid Core Derived Peptides.** Panels A), B), C), D) and E) refer to peptides corresponding to regions 3-13, 47-57, 79-95, 105-120 and 135-141, respectively. The blue curves correspond to wild type peptide whereas the red curves correspond to the sequences with phosphor-mimetic substitutions. Free energy minimum in the region shaded yellow corresponds to a stable self-assembled state. Minima in the region shaded blue refers to a fully mixed state of the peptides.

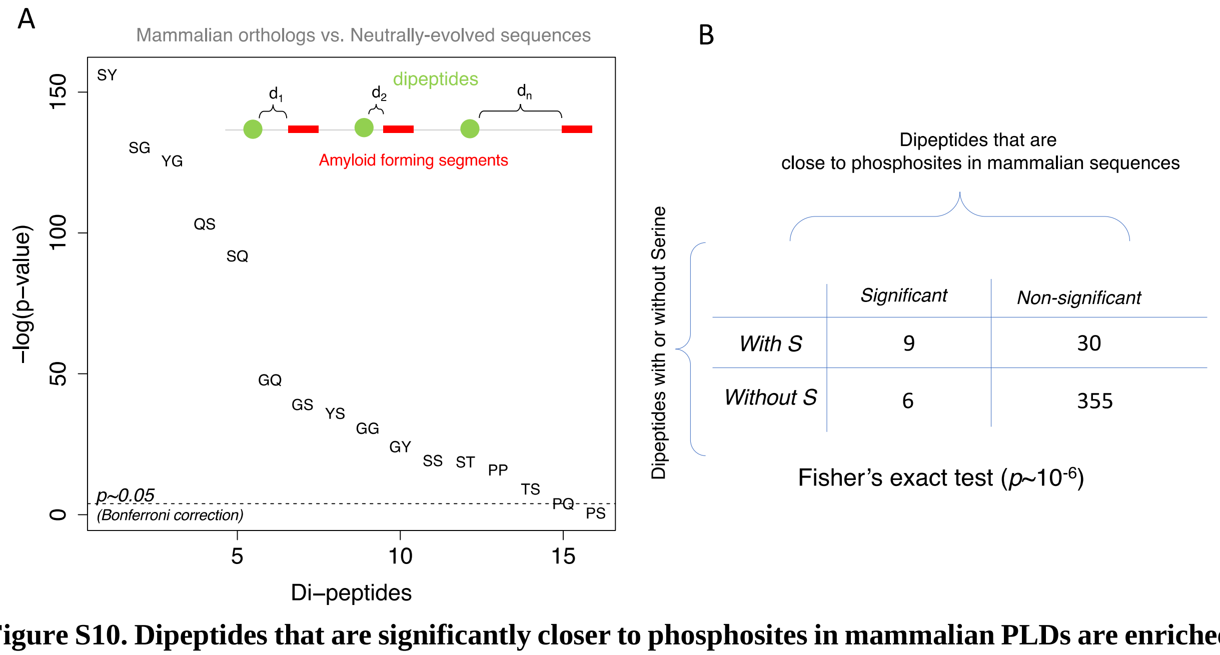


| | Significant | Non-significant |
|---|---|---|
| With S | 9 | 30 |
| Without S | 6 | 355 |

**Figure S10. Dipeptides that are significantly closer to phosphosites in mammalian PLDs are enriched in serine residues.** A) –log(p-value) versus the rank of dipeptides in their proximity to the potential phosphorylation sites. Here, p-values are calculated from the one-sided wilcoxon rank-sum test with the alternative hypothesis that dipeptides are closer to potential phosphosites. We applied the Bonferroni correction by dividing the unadjusted p-value (0.05) by the total number of possible dipeptides (=400). B) The contingency table and the result of Fisher's exact test to investigate whether dipeptides that are closer to potential phosphorylation sites are enriched in serine residues.

#### Supplementary Tables

**Table S1. Amino Acid Parameters used for residue-level coarse-graining using the HPS model.**

| Type | Charge | Size ($\sigma$) | Hydrophobicity (H) |
|---|---|---|---|
| ALA | 0 | 5.04 | 0.730 |
| ARG | 1 | 6.56 | 0 |
| ASN | 0 | 5.68 | 0.432 |
| ASP | -1 | 5.58 | 0.378 |
| CYS | 0 | 5.48 | 0.595 |
| GLN | 0 | 6.02 | 0.514 |
| GLU | -1 | 5.92 | 0.459 |
| GLY | 0 | 4.5 | 0.65 |
| HIS | 0.5 | 6.08 | 0.514 |
| ILE | 0 | 6.18 | 0.973 |
| LEU | 0 | 6.18 | 0.9730 |
| LYS | 1 | 6.36 | 0.514 |
| MET | 0 | 6.18 | 0.838 |
| PHE | 0 | 6.36 | 1 |
| PRO | 0 | 5.56 | 1 |
| SER | 0 | 5.18 | 0.595 |
| THR | 0 | 5.62 | 0.676 |
| TRP | 0 | 6.78 | 0.946 |
| TYR | 0 | 6.46 | 0.865 |
| VAL | 0 | 5.86 | 0.892 |

**Supplementary Table S2. Human FUS Sequence**

| |
|---|
| >sp\|P35637\|FUS_HUMAN RNA-binding protein FUS OS=Homo sapiens OX=9606 GN=FUS PE=1 SV=1<br>MASNDYTQQATQSYGAYPTQPGQGYSQQSSQPYGQQSYSGYSQSTDTSGYGQSSYSS<br>YGQSQNTGYGTQSTPQGYGSTGGYGSSQSSQSSYGQQSSYPGYGQQPAPSSTSGSYGS<br>SSQSSSYGQPQSGSYSQQPSYGGQQQSYGQQQSYNPPQGYGQQNQYNSSSGGGGG<br>GGGGNYGQDQSSMSSGGGSGGGYGNQDQSGGGGSGGYGQQDRGGRGRGGSGGGG<br>GGGGGGYNRSSGGYEPRGRGGGRGGRGGMGGSDRGGFNKFGGPRDQGSRHDSEQD<br>NSDNNTIFVQGLGENVTIESVADYFKQIGIIKTNKKTGQPMINLYTDRETGKLKGEATV<br>SFDDPPSAKAAIDWFDGKEFSGNPIKVSFATRRADFNRGGGNGRGGRGRGGPMGRGG<br>YGGGGSGGGGRGGFPSGGGGGGGQQRAGDWKCPNPTCENMNFSWRNECNQCKAP<br>KPDGPGGGPGGSHMGGNYGDDRRGGRGGYDRGGYRGRGGDRGGFRGGRGGGDRG<br>GFGPGKMDSRGEHRQDRRERPY |

**Supplementary Table S3. Human FUS Phosphosites**

| | |
|---|---|
| | S3,T7,T11,T19,S26,S30,S37,S42,S54,S57,S61,T68,T71,S77,T78,S84,S86,S87,S95,S96,S108,S109,S110,S112,S115,S117,S127,S129,S131,S135,S142,S148 |

**Supplementary Table S4.**
**Amyloid-core derived peptide sequences used in atomistic simulations**

| Amyloid Core | Region of FUS sequence | Sequence | Phosphosites | Phosphomimetic Variant |
|---|---|---|---|---|
| C1 | 3-13 | **S**NDY**T**QQA**T**QS | S3,T7,T11 | **E**NDY**E**QQA**E**QS |
| C3 | 47-57 | TSGYGQS**S**YS**S** | S54,S57 | TSGTGQS**E**YS**E** |
| C4 | 79-95 | GGYGS**S**Q**SS**QSSYGQQ**S** | S84,S86,S87,S95 | GGYGS**E**Q**EE**QSSYGQQ**E** |
| C5 | 105-120 | AP**STS**G**S**YG**S**S**S**QSS | S108,T109,S110,S112,S115,S117 | AP**EEE**G**E**YG**E**S**E**QSS |
| C7 | 135-141 | **S**YGGQQQ | S135 | EYGGQQQ |
| polyG | -- | GGGGGGGGG | -- | -- |
